# Loss of Lamin A leads to the nuclear translocation of AGO2 and compromised RNA interference

**DOI:** 10.1101/2023.06.05.543674

**Authors:** Vivian Lobo, Iwona Nowak, Carola Fernandez, Ana Iris Correa Muler, Jakub O. Westholm, Hsiang-Chi Huang, Ivo Fabrik, Hang Thuy Huynh, Evgeniia Shcherbinina, Melis Poyraz, Anetta Härtlova, Daniel Benhalevy, Davide Angeletti, Aishe A. Sarshad

## Abstract

In mammals, RNA interference (RNAi) was historically studied as a cytoplasmic event; however, in the last decade, a growing number of reports convincingly show the nuclear localization of the Argonaute (AGO) proteins. Nevertheless, the extent of nuclear RNAi and its implication in biological mechanisms remain to be elucidated. We found that reduced Lamin A levels significantly induce nuclear influx of AGO2 in SHSY5Y neuroblastoma and A375 melanoma cancer cell lines, which normally have no nuclear AGO2. Lamin A KO manifested a more pronounced effect in SHSY5Y cells compared to A375 cells, evident by changes in cell morphology, increased cell proliferation, and oncogenic miRNA expression. Furthermore, in SHSY5Y cells, AGO fPAR-CLIP in Lamin A KO cells revealed significantly reduced activity of RNAi. Further exploration of the nuclear AGO interactome by mass spectrometry indicated that AGO2 is in complex with FAM120A, an RNA-binding protein and known interactor of AGO2. By performing FAM120A fPAR-CLIP, we discovered that FAM120A co-binds AGO targets and that this competition reduces the activity of RNAi. Therefore, loss of Lamin A triggers nuclear AGO2 translocation, RNAi impairment, and selective upregulation of oncogenic miRNAs, facilitating cancer cell proliferation.

## INTRODUCTION

The eukaryotic genome is enclosed by the nuclear envelope (NE), which is a highly regulated barrier that separates the nucleus from the cytoplasm [1]. One of the major components of the NE are the nuclear Lamins [2], encoded by three highly conserved Lamin genes – *LMNA*, *LMNB1* and *LMNB2* [3]. Alternative splicing of *LMNA* gene results in the production of four different A-type Lamin proteins, where Lamin A and Lamin C are the predominant isoforms [4]. While *LMNB1* and *LMNB2* encoded proteins (B-type Lamins) are ubiquitously expressed, A-type Lamin expression is limited to differentiated cells [3].

A and B type Lamins belong to the type V intermediate filaments protein family and assemble into higher order structures forming the nuclear lamina – a main component of the NE and an important platform for information exchange between the nucleus and cytoplasm [5]. Accordingly, Lamin proteins highly affect gene expression. The Lamins are closely associated with Lamin-associated domains (LADs), genomic regions characterized by densely packed chromatin and transcriptional inactivation [6–7]. Therefore, precise control of the nuclear lamina composition is crucial for safeguarding cellular identity by keeping proper chromatin organization and cellular morphology [8–9].

Gene expression is also regulated post-transcriptionally, among others by RNA interference (RNAi), executed by miRNA-loaded Argonaute (AGO) proteins [10–11]. In mammals, miRNAs matured by DROSHA and DICER enzymatic processing, are loaded into one of four AGO proteins, comprising the core RNA induced silencing complex (RISC). Based on sequence complementation AGO:miRNA RISC target mRNAs [12]. RISC induces RNAi by recruiting key mediating factors, such as TNRC6A-C scaffolding proteins, which bridge the RISC complex with the CCR4-NOT deadenylation complex. In metazoans, the process of RNAi is generally accepted as a cytoplasmic event [13–14], but numerous reports have documented the nuclear localization of AGO proteins [15–19]. Conditions such as cell confluence, oxidative stress, quiescence, and senescence have been found to trigger AGO2 nuclear influx [20–23]. The mechanism of AGO2 nuclear entry is so far not completely delineated, but there is evidence that the nuclear localization of AGO2 may directly be regulated by its interacting partner TNRC6 [20]. Additionally, nuclear import of AGO:miRNA complexes has also been linked to Importin 8 in HeLa cells and more recently to FAM172A in mouse embryonic fibroblasts [15,24]. In the nucleus, AGO proteins have been suggested to regulate several processes aside from RNAi, including chromatin remodeling, transcriptional regulation, alternative splicing, genome integrity, and DNA repair in cancer cells [15-18; 25-26]. Therefore, AGOs may promote the regulation of several biological processes in a context-dependent manner, based on the cellular environment and RNA:protein interactors.

Since RNAi is a pivotal process for post-transcriptional gene silencing, its activity is tightly regulated to ensure homeostasis. Cancer progression is associated with the impairment of RNAi and global miRNA downregulation [27]. Apart from inhibition of miRNA biogenesis pathway and decreased AGO protein expression, malignant cells are characterized by the reduced activity of AGO:miRNA [28–31]. One mode for negative regulation of RNAi in cancer is the post-translational modifications of AGO proteins catalyzed by oncogenic kinases such as EGFR, Akt, and Src [28–31]. Moreover, numerous oncogenic RNA binding proteins (RBPs), such as HuR, HuD, and IGF2, were shown to stabilize pro-proliferative transcripts by competitive AGO binding [32–34]. Notably, a recent report described the nuclear influx of AGO2 in overconfluent cancer cell cultures, as well as in malignant colon cancer tissues [20]. Interestingly, this was accompanied by RNAi inhibition [20].

Yet, the underlying molecular mechanisms of AGOs’ nuclear translocation and putative nuclear RNAi-mediated gene regulation in cancer cells remain to be fully explored. Several groups have investigated nuclear AGO2 transcript occupancy and reported substantial occupancy of AGO2 on intronic sites within pre-mRNAs [18, 25, 35]. In stem cells, nuclear RNAi reflected cytoplasmic RNAi, but was significantly more potent [18]. However, most of the literature does not address the level of nuclear AGO-mediated gene silencing.

Since the nuclear envelope is a crucial link in a chain of dialogue between the cytoplasm and the nucleus, its composition is bound to influence protein transportation. However, this is not extensively characterized. Further, the cellular state and factors leading to increased nuclear AGO2 accumulation and regulation are not yet fully elucidated. Despite that the first observation of nuclear AGO2 dates to 2008 [36], the extent of the phenomenon, and AGOs nuclear activity and biological function remain to be fully explored. Herein, we aimed to investigate the effect of the composition of the nuclear lamina on AGO2 nuclear translocation.

We identified that loss of Lamin A expression triggers AGO nuclear translocation, and upregulation of pro-oncogenic miRNAs in cancer cells. Furthermore, loss of Lamin A leads to oxidative stress, a condition which activates FAM120A, a known interactor of AGO2. We discovered that FAM120A co-binds AGO targets and that this competition significantly reduces the activity of RNAi upon loss of Lamin A.

## MATERIAL AND METHODS

### Cell culture

HeLa ovarian adenocarcinoma cells (ATCC, CCL-2), MCF7 breast cancer cells (ATCC, HTB-22), CUT09 and STE01 lung cancer cells (kind gift from Ruth Palmer and Bengt Hallberg), Molm13 acute myeloid leukemia cells (DMSZ, ACC 554), Granta-519 B cell lymphoma cells (DMSZ, ACC 342), HEK293 (ATCC, CRL-1573) and HEK293T (ATCC, ACS-4500) embryonic kidney cells, SHSY5Y neuroblastoma cells (ATCC, CRL-2266), HAP1 chronic myeloid leukemia cells (kind gift from Maria Falkenberg), U2OS osteosarcoma cells (ATCC, HTB-96) and A375 melanoma cells (ATCC, CRL-1619) were cultured at 37°C, with 5% CO_2_. HeLa, MCF7, HEK293, HEK293T, U2OS and A375 cells were cultured in DMEM media (11995065, Gibco) supplemented with 10% Fetal Bovine Serum (FBS) (10270106, Gibco) and 1% Penicillin-Streptomycin (P/S) solution (15140122, Gibco). CUT09, STE01, Molm13, Granta-519 and SHSY5Y cells were cultured in RPMI media (61870010, Gibco) complemented with 10% FBS and 1% P/S solution. HAP1 cells were cultured in IMDM media (12440053, Gibco) supplemented with 10% FBS and 1% P/S solution. A375 and SHSY5Y Lamin A knock out cells were cultured with respective complete media complemented with 2 μg/ml puromycin (P9620, Sigma Aldrich) and 350 μg/ml G418 (G8168, Sigma Aldrich).

### Biochemical fractionation

Cells were fractionated as described in [37] with minor modifications. Briefly, cells were resuspended in Hypotonic Lysis Buffer (HLB) buffer [10 mM Tris-HCl, pH 7.5; 10 mM NaCl; 3 mM MgCl2; 0.3% NP-40 (vol/vol); and 10% glycerol (vol/vol)], supplemented with protease and phosphatase inhibitor cocktail (78445, ThermoFisher) by gentle pipetting and left on ice for 5 min. The nuclei were separated by centrifugation at 200 *g* for 2 min and the supernatant was collected as the cytoplasmic lysate. The pellet, containing the nuclear fraction, was further washed three times in HLB buffer and each time collected by centrifugation at 200 *g* for 2 min. The washed nuclear pellet was resuspended with equal volumes of Nuclear Lysis Buffer (NLB) buffer [20 mM Tris-HCl, pH 7.5; 150 mM KCl; 3 mM MgCl2; 0.3% NP-40 (vol/ vol), and 10% glycerol (vol/vol)], supplemented with protease and phosphatase inhibitor cocktail. The nuclear lysate was sonicated twice, 10 s on and 30 s off, at 60% amplitude (Sonics, VCX130). Both cytoplasmic and nuclear lysates were further cleared by centrifugation at 21,000 *g* for 15 min.

### Western blot analysis

Total protein lysates were prepared by suspending the cell pellets in NP-40 lysis buffer [20 mM Tris-HCl, pH 7.5; 150 mM NaCl; 2 mM EDTA; 1% (v/v) NP-40] and incubating on ice for 20 min. Following, the samples were centrifuged at 21,000 *g* for 20 min and the supernatant collected as the total protein lysate. Cytoplasmic and nuclear lysates were prepared as described above. Protein concentration for the total and cytoplasmic lysates were quantified by Bradford Reagent (B6916, Sigma Aldrich) and 10-20 μg of protein was used for western blot experiments. The nuclear lysates were loaded at 1:1 volume ratio with the cytoplasmic fraction, on to 8%-12% Bis-Tris-HCl PAGE gels. Following electrophoresis, the proteins were transferred onto Nitrocellulose membrane (1060000, Cytiva) and the blots stained with Ponceau S, washed, and blocked with skimmed milk (70166, Sigma). Membranes were incubated overnight with either of the following antibodies indicated in table below. Following, blots were washed and incubated with HRP conjugated anti-rabbit IgG (NA934, Cytiva) or anti-mouse IgG (NA931, Cytiva). The signal was detected by chemiluminescence using Thermo Scientific SuperSignal™ West Dura Extended Duration Substrate (34076, ThermoFisher).

| Antibody | Company | Dilution |
| --- | --- | --- |
| anti-AGO1 | 5053, CST | 1:1000 |
| anti-AGO2 | ab32381 or ab186733, Abcam | 1:5000 |
| anti-AGO3 | 5054, CST | 1:1000 |
| anti-Lamin B1 | ab16048, Abcam | 1:10,000 |
| anti-Histone H3 | 4499, CST | 1:10,000 |
| anti-Lamin A | sc-518013, Santacruz | 1:1000 |
| anti-Lamin A/C | sc-376248HRP, Santacruz | 1:1000 |
| Streptavidin-HRP | S911, Molecular Probes | 1:4000 |
| anti-V5 | R960-25, Invitrogen | 1:4000 |
| anti-FAM120A | ab156695, Abcam | 1:2000 |
| anti-FAM120A | ABIN7451012, Antibodies online | 1:1000 |
| anti-PABPC | sc-32318, Santacruz | 1:1000 |
| anti-Calreticulin | sc-373863HRP, Santacruz | 1:1000 |
| anti-TNRC6A | A302-329A, Bethyl Laboratories | 1:1000 |

### siRNAs transfection

siRNAs targeting *LMNA* (4392420 and 4390824, assay IDs: s530951 and s8221) and *LMNB1* (4392420, assay IDs: s8224 and s8225), as well as negative control scramble oligonucleotide (4390843, 4390846), without complementarity to sequences in the human transcriptome, were purchased from ThermoFisher. Cells were seeded one day prior to the siRNA transfections. siRNAs, scramble oligonucleotide and Lipofectamine RNAiMax (13778150, ThermoFisher) were diluted in Opti-MEM reduced media (31985047, Gibco) and added to the cell culture at a final concentration of 7.5 nM. 24 h following the transfection, the media was changed to fresh complete media. After another 24 h of culture, cells were harvested for downstream analysis. Silencing efficiency was assessed by western blot.

### CRISPR-Cas9 gene editing

Lamin A knockout cells were generated in A375, SHSY5Y and HeLa cells using PITCH-CRISPR-Cas9-microhomology-mediated end-joining [38]. Briefly, two all-in-one CRISPR-Cas9 vectors were generated from pX330A-1×2 (58766, Addgene) and pX330S-2-PITCh (63670, Addgene), expressing Cas9 nuclease and two gRNAs (see below for sequences) together with the CRIS-PITCh vector pX330S-2-PITCh (63670, Addgene), harboring the Lamin A microhomologies and GFP-Puro or GFP-Neo/Kan insertions, were transiently transfected into A375 and SHSY5Y using JetOptimus transfection reagent (117-01, Polyplus) according to the manufacturer’s instructions. Single clones undergoing recombination were selected by complementing culture media with Puromycin at a concentration of 2 μg/ml and G418 at a concentration of 350 μg/ml for 10 days. Homozygous Lamin A knockout was identified by PCR and western blot screening. Genomic DNA was extracted using PureLink™ Genomic DNA (K182002, ThermoFisher) following the manufacture’s protocol. PCR was performed using OneTaq polymerase (M0480S, New England Biolabs) following the manufacture’s protocol. The primer sequences were as follows: primer 232: GCAGATCAAGCGCCAGAATG; primer 233: GCTTCATGTGGTCGGGGTAA; primer 235: GGGACACTGTCAAGCTCTCC;; primer 281: TTACCCCGACCACATGAAGC; β-Actin Fw: GTCGTCGACAACGGCTCCGGCATGTG, Rv: CATTGTAGAAGGTGTGGTGCCAGAT.

The gRNA sequences targeting *LMNA* Exon 9 locus and a generic PITCh-gRNA were as follows: *LMNA* sgRNA-Ex 9 Fw1: aaacCTACCGACCTGGTGTGGAAGC; *LMNA* sgRNA-Ex 9 Rv1: CACCGCTTCCACACCAGGTCGGTAG; *LMNA* sgRNA-Ex 9 Fw2: CACCGAGTTGATGAGAGCCGTACGC; *LMNA* sgRNA-Ex 9 Rv 2: aaacGCGTACGGCTCTCATCAACTC)

### Immunofluorescence staining

Cells were plated at a density of 60,000 cells/cm^2^ on glass coverslips inside 12-well dishes. Next day, cells were washed once in 1x PBS and fixed with 4% paraformaldehyde (1267, Solveco) for 15 min at RT. Following, cells were washed with 1x PBS and permeabilized in 0.5% Triton X-100 for 10 min at RT. Subsequently, cells were blocked with 5% BSA (BP9703-100, Fisher Bioreagents) in 1x PBS for 1 h and stained with primary antibodies indicated in the table below. The cells were washed three times in 1x PBS and stained with 1:400 Phalloidin (B3475, Invitrogen), to visualize F-actin, or secondary antibodies indicated in the table below for 1 h at RT. To image the Endoplasmic Reticulum after fixing with paraformaldehyde, cells were incubated with 1 μM of ER-Tracker Red dye (E34250, Invitrogen) for 30 minutes at 37°C. To visualize the DNA within the nucleus, cells were stained with 300 nM 4’,6-diamidino-2-phenylindole (DAPI; D1306, Invitrogen) for 5 min at RT. Slides were mounted using ProLong Diamond Antifade Mountant (P36961, Invitrogen). Confocal images were taken on a Zeiss LSM780 and the images analyzed using ImageJ software.

| Antibody | Company | Dilution |
| --- | --- | --- |
| anti-Lamin B1 | ab16048, Abcam | 1:500 |
| anti-Pol II | sc-56767 Santa Cruz | 1:200 |
| anti-Lamin A | sc-518013, Santacruz | 1:400 |
| anti-Histone H3 | 4499, CST | 1:400 |
| anti-V5 | R960-25, Invitrogen | 1:400 |
| anti-rabbit Alexa 488 | A11008, Invitrogen | 1:2000 |
| anti-mouse Alexa 594 | A11005, Invitrogen | 1:2000 |

### Cell proliferation assay

For cell proliferation assays, A375 and SHSY5Y (WT and Lamin A KO) cells were seeded in 6-well plates at density of 3,125 cells/cm^2^ and counted with LUNA-II™ Automated Cell Counter (Logos Biosystems) every 24 h for 6 days. The cells were grown in their respective complete media, which was changed every two days. The experiment was performed in four biological replicates and, additionally, each included two technical replicates. Exponential (Malthusian) growth proliferation assay line was fitted, and doubling time calculated using GraphPad Prism (Version 9.5.1).

### Flow cytometry for cell cycle analysis

A375 and SHSY5Y (WT and Lamin A KO) cells were seeded in 6-well plates at a density of 10,500 cells/cm^2^ and incubated at 37°C overnight. For cell cycle analysis, 50,000 cells were collected and fixed with ice-cold 70% (v/v) ethanol at -20°C overnight. The cells were pelleted at 300 *g* for 5 min and the pellet was washed once with PBS at 300 *g* for 5 min and subsequently suspended in PI solution (50 μg/ml Propidium iodide (81845, Sigma Aldrich), 0.1 mg/ml RNase A (EN0531, ThermoFisher), 0.05% Triton X-100 in 1x PBS) for 40 min at 37°C. Cells were then washed with 1x PBS and processed for cell cycle analysis on BD-LSRII LSRII (BD Bioscience). Data was analyzed using FlowJo (Treestar Inc.).

### Plasmid transfection

Cells were transfected with the following expression plasmids: eGFP-NLS (67652, Addgene), pCDH-CMV-hLamin_A-IRES-copGFP-EF1-puro (132773, Addgene), using JetOptimus transfection reagent (117-01, Polyplus) or Lipofectamine 2000 (11668027, Invitrogen) according to manufacturers’ instructions for 24-48 h. In case of transfections with pCDH-CMV-hLamin_A-IRES-copGFP-EF1-puro plasmid cells were selected with 2 μg/ml puromycin for 24 h prior to collection. Transfection efficiency was assessed by immunofluorescence staining and/or western blot.

### RNA isolation, reverse transcription, and qPCR

Total RNA was extracted using Quick-RNA Miniprep Kit (BioSite-R1055, Zymo Research). RNA concentration was assessed with Implen N60 nanophotometer. Following, 500 ng of RNA was reverse transcribed using iScript™ cDNA Synthesis Kit (1708891, Biorad). For qPCR reactions, cDNA was diluted 50 times. qPCR was performed in XT96 thermocycler with iTaq™ Universal SYBR® Green Supermix (1725124, Biorad) using thermocycling conditions recommended by the manufacturer. The sequences of the primers were as follows: 18S rRNA: Fw: CCCTCCAATGGATCCTCGTT, Rv: AGAAACGGCTACCACATCCA, *AGO2:* Fw: TGAGCTTGCAGGCGTTACAC, Rv: CAAGAGGGTTAGAGCAGCCTT; *LMNA*: Fw: AGACCCTTGACTCAGTAGCC, Rv: AGCCTCCAGGTCCTTCA.

### Generation of V5-APEX2-SENP2 expressing cells

V5-APEX2-SENP2 expression was introduced into A375 Lamin A KO and WT cells by lentiviral transduction according to the protocol described by Tandon and colleagues [39]. Cells were co-transfected with plasmids encoding a mixture of viral packaging proteins VSV-G (12259, Addgene), viral backbone psPAX2 plasimd (12260, Addgene) and V5-APEX2-SENP2 (129276, Addgene) at ratio 3:2:4, where the final plasmid concentration in the media was 2.25 μl/ml. Cells were kept in selection media [complete media supplemented with 10 μg/ml Blasticidin (A1113903, Gibco)] for 10 days, when control, none transduced, cells died. Following, cells were cultured in complete media for respective parental cell lines.

### Proximity-dependent biotinylation

Cells were grown in 15 cm cell culture dishes until they reached approximately 50% confluency. Following, biotinylation of proteins proximal to V5-APEX2-SENP2 was performed according to previously published protocol [40] with a minor modification where Biotin-Phenol (SML2135, Sigma-Aldrich) was added to the media at a final concentration of 1 mM and the incubation with Biotin-Phenol was extended to 40 min.

### Streptavidin affinity pull down assay

Following the proximity-dependent biotinylation procedure, cells were lysed in RIPA buffer [50 mM Tris-HCl, 150 mM NaCl, 0.1% (w/v) SDS, 0.5% (w/v) sodium deoxycholate and 1% (v/v) Triton X-100] complemented with 10 nM sodium azide (S2002, Sigma-Aldrich), 10 mM sodium ascorbate (A4034, Sigma-Aldrich) and 5 nM of Trolox (238813, Sigma-Aldrich) and protease inhibitor (78439, ThermoFisher). Subsequently, the protein concentration was estimated using Pierce reagent (22660, ThermoFisher), and 250 µg of protein were subjected to streptavidin affinity precipitation using 20 μl streptavidin-coupled magnetic beads (88817, ThermoFisher) at 4°C overnight, under constant rotation, for enrichment of biotinylated proteins. After the incubation, beads were washed with a series of buffers: 2 times with RIPA buffer, once with 1 M KCl (10735874, ThermoFisher), once with 0.1 M Na_2_CO_3_ (230952, Sigma-Aldrich), once with 2 M Urea (U6504, Sigma-Aldrich) in 10 mM Tris-HCl-HCl (15567027, Invitrogen™), freshly prepared, and twice with RIPA buffer. Following the washes, beads were incubated for 10 min at 95°C in 60 μl of 3x Laemmli SDS Sample Buffer (J61337.AC, ThermoFisher) supplemented with 2 nM biotin (B4501, Sigma-Aldrich), to elute bound proteins for subsequent western blot analysis.

### Immunoprecipitation assays and AGO protein affinity purification with T6B peptide

For identification of Lamin A and AGO-interacting protein partners, respective proteins were pulled down using either anti-Lamin A antibody or Flag-tagged T6B peptide for AGO1-4 isolation [41]. Lamin A was immunoprecipitated from 2 mg of total protein lysate in A375 WT and Lamin A KO cells with 3 µg of antibody. AGO1-4 were pulled down from nuclear or cytoplasmic lysates. Volumes corresponding to 3 mg of cytoplasmic fraction was used together with 400 μg of T6B peptide. Dynabead Protein G beads (10004D, ThermoFisher) and anti-Flag M2 beads (M8823, Millipore) were conjugated with either anti-Lamin A antibody or T6B peptide for 4 h, washed and incubated with protein lysates. Next, beads were washed with NP-40 buffer. Lamin A and T6B bound proteins were eluted by incubation with 0.2 M Glycine, pH 2.5 by gentle shaking for 15 min followed by neutralization of the eluate using 1 M Tris-HCl, pH 8. The pull-down efficiency was confirmed by western blot and, in case of AGO1-4 and Lamin A pull down, the eluate submitted for mass spectrometry analysis at Proteomics Core Facility (University of Gothenburg, Sweden).

### Mass spectrometry

Samples eluted after AGO1-4 proteins and Lamin A affinity purification were processed using modified filter-aided sample preparation (FASP) method [42]. Briefly, samples were reduced in 100 mM dithiothreitol (DTT) at 60°C for 30 min, transferred to Microcon-30kDa Centrifugal Filter Units (MRCF0R030, Merck) and washed several times with 8 M urea and once with digestion buffer (DB; 50 mM TEAB, 0.5% sodium deoxycholate (SDC)) prior to alkylation (10 mM methyl methanethiosulfonate (MMTS) in DB for 30 min at RT. Samples were digested with 0,3 µg Pierce MS grade Trypsin (90057, ThermoFisher) at 37°C overnight and an additional portion of trypsin was added and incubated for another two h. Peptides were collected by centrifugation.

Following, digested peptide samples were labelled using TMT 16-plex isobaric mass tagging reagents (90063, ThermoFisher) for relative quantification. The samples were combined into one TMT-set and SDC was removed by acidification with 10% TFA. The TMT-set was further purified using High Protein and Peptide Recovery Detergent Removal Spin Column (88305, ThermoFisher) and Pierce peptide desalting spin columns (89852, ThermoFisher) according to the manufacturer’s instructions, prior to basic reversed-phase chromatography (bRP-LC) fractionation. Peptide separation was performed using a Dionex Ultimate 3000 UPLC system (ThermoFisher) and a reversed-phase XBridge BEH C18 column (3,5 μm, 3.0×150 mm, 186008164, Waters Corporation) with a gradient from 3% to 100% acetonitrile in 10 mM ammonium formate at pH 10.00 over 23 min at a flow of 400 µL/min. The 40 fractions were concatenated into 20 fractions.

The dried samples were reconstituted in 3% acetonitrile and 0,2% formic acid. The TMT fractions in were analyzed on an Orbitrap Lumos Tribrid mass spectrometer interfaced and an Easy-nLC1200 liquid chromatography system (ThermoFisher). Peptides were trapped on an Acclaim Pepmap 100 C18 trap column (100 μm x 2 cm, particle size 5 μm, 164199, ThermoFisher) and separated on an in-house packed analytical column i.d. 75 μm, particle size 3 μm, Reprosil-Pur C18, Dr. Maisch, length 35 cm using a gradient from 3% to 80% acetonitrile in 0.2% formic acid over 85 min at a flow of 300 nL/min. Precursor ion mass spectra were acquired at 120,000 resolution and MS/MS analysis was performed in data-dependent mode where CID spectra of the most intense precursor ions were recorded in the ion trap at collision energy setting of 35. Precursors were isolated in the quadrupole with a 0,7 m/z isolation window, charge states 2 to 7 were selected for fragmentation, dynamic exclusion was set to 45 s and 10 ppm. MS3 spectra for reporter ion quantitation were recorded at 50,000 resolution, multi-notch isolation with 10 notches and HCD fragmentation collision energy of 55.

Lamin A affinity purification samples were analyzed by label free approach on an Orbitrap Exploris 480 mass spectrometer interfaced with an Easy-nLC1200 liquid chromatography system (ThermoFisher) in trap column configuration (see above). Peptides were separated using a gradient from 5% to 45% acetonitrile in 0.2% formic acid over 48 min at a flow of 300 nL/min. Precursor ion mass spectra were acquired at 120,000 resolution and MS/MS analysis was performed in data-dependent mode with HCD collision energy setting of 30. Precursors were isolated in the quadrupole with a 0.7 m/z isolation window, charge states 2 to 6 were selected for fragmentation, dynamic exclusion was set to 30 s and 10 ppm.

Following, the data files were merged for identification and relative quantification using Proteome Discoverer version 2.4 (ThermoFisher). The data matching was against Homo Sapiens (Swissprot) using Mascot version 2.5.1 (Matrix Science) as a search engine. The precursor mass tolerance was set to 5 ppm and fragment mass tolerance to 0,6 Da. Tryptic peptides were accepted with one missed cleavage, variable modifications of methionine oxidation and fixed cysteine alkylation, TMT pro modifications of N-terminal and lysine were selected. Percolator was used for PSM validation with the strict FDR threshold of 1%. Identified proteins were filtered at 1% FDR at protein level. For relative quantification reporter ions were identified in the MS3 HCD spectra with 3 mmu mass tolerance. Only the quantitative results for the unique peptide sequences with the minimum SPS match % of 50 and the average S/N above 10 were considered for the protein quantification. Label free quantification was done in MaxQuant 2.1.3.0 [43] by searching against Homo sapiens database (Uniprot). The precursor mass tolerance was set to 4.5 ppm and fragment mass tolerance to 20 ppm. Tryptic peptides were accepted with two missed cleavages, variable modifications of methionine oxidation and N-term protein acetylation and fixed cysteine alkylation. FDR threshold for identification was set to 1%. Proteins were quantified by MaxLFQ algorithm [44].

### Mass spectrometry data interpretation

TMT reporter intensities from AGO affinity purification samples were normalized in Perseus ver. 2.0.7.0 [45] so that medians of log2-ratios between Lamin KO and WT for each cell line were 0. Obtained intensities were log2-transformed, tested by Student’s T-test and proteins passing 1,5 fold change and p<0.075 were considered as having significantly enhanced interaction with AGO proteins in Lamin A KO. MaxLFQ intensities from Lamin A affinity purification samples were log2-transformed and tested as stated above. Enrichment of Gene Ontology Biological Process terms for significant hits from each cell line was determined by DAVID web-based tool [46]. The interaction network was constructed in Cytoscape ver. 3.8.2 [47] based on protein-protein interaction data from STRING ver. 11.5 [48] for significant proteins common for both cell lines and adding AGO2 as IP bait.

### RNA sequencing

Total RNA was extracted using Quick-RNA Miniprep Kit. Following, the concentration and quality of the RNA was analyzed using Agilent 2200 TapeStation System. RNA samples with RNA Integrity Number higher than 8 were sent to SNP&SEQ Technology Platform (NGI Uppsala, Sweden). Libraries was prepared from 300 ng RNA using the Illumina Stranded Total RNA library preparation kit, including Ribo-Zero Plus treatment (20040525/20040529, Illumina Inc.) according to manufacturer’s instructions. For indexing Unique Dual Indexes (20040553/20040554, Illumina Inc.) were used. Sequencing was carried out with NovaSeq 6000 system using paired-end 150 bp read length, S4 flowcell and v1.5 sequencing chemistry. As a control sequencing library for the phage PhiX was included and a 1% spike-in in the sequencing run. RNAseq data were preprocessed using the RNAseq nf-core pipeline [49]. Differential expression analysis was done using DEseq2 [50], on genes with at least 10 reads in at least 3 samples. Genes with FDR adjusted p-value < 0.01 and absolute log2 fold change > 0.5 were considered differentially expressed. Hypergeometric test, implemented in TopGO, were used to look for enriched Gene Ontology annotation among the differentially expressed genes. The fraction of reads mapping to introns and other genomic regions was calculated using ReSQC [51].

### miRNA sequencing

AGO proteins were immunoprecipitated from 1.5 mg of protein using Flag-tagged T6B peptide. As a negative control immunoprecipitation with empty beads were used. Following, RNA was recovered from the beads using TRIzol reagent (15596026, Invitrogen) according to the manufactures instructions and small RNA libraries were produced as previously described [52] with few modifications. Briefly, 3’ adapters with 5’-adenylated RNA adapter (see 3’ adapters in table below) were ligated to the recovered small RNAs using Rnl2(1-249)K227Q RNA ligase (M0351, New England Biolabs) according to the manufacturer’s instructions at 4°C overnight with shaking. Ligated RNA was pooled and purified using oligo clean and concentrate kit (D4060, ZYMO Research). Following, the RNA was subjected to 5’ adapter ligation with a 5’ chimeric DNA-RNA adapter (5’aminolinker-GTTCAGAGTTCTACAGTCCGACGATCrNrNrNrN) using RNA ligase (EL0021, Thermo Fisher Scientific) at 37°C for 1 hour. Next, the RNA was purified using oligo clean and concentrate kit and reverse transcribed using SuperScript® IV (18090010, Thermo Fisher Scientific) as per the protocol provided by manufacturer using RT primer (GCCTTGGCACCCGAGAATTCCA). The cDNA was amplified using Platinum Taq DNA Polymerase (10966034, Thermo Fisher Scientific), according to the manufacturer’s instructions using 5’-medium PCR primer (CTCTACACGTTCAGAGTTCTACAGTCC) and 3’ medium PCR primer (CCTGGAGTTCCTTGGCACCCGAGAATT) for 6 cycles. Then the PCR product was purified using the oligo clean and concentrate kit, eluted with 32 µl of nuclease free water, and size selected (74-88 bp) using 3% agarose Pippin Prep (CSD3010, Sage Science). Following size selection, a second round of (X cycle) PCR was performed using the same polymerase, a 5’-long PCR primer: (AATGATACGGCGACCACCGAGATCTACACGTTCAGAGTTCTACAGTCCGA), and 3’ indexed primer (see 3’ index primers in table below), Libraries were sequenced on an Illumina NovaSeq6000. Bcl files were converted to fastq files using bcl2fastq. Adapters were trimmed using cutadapt v 2.4. and reads were mapped to the human miRNAs using bowtie2 [53].

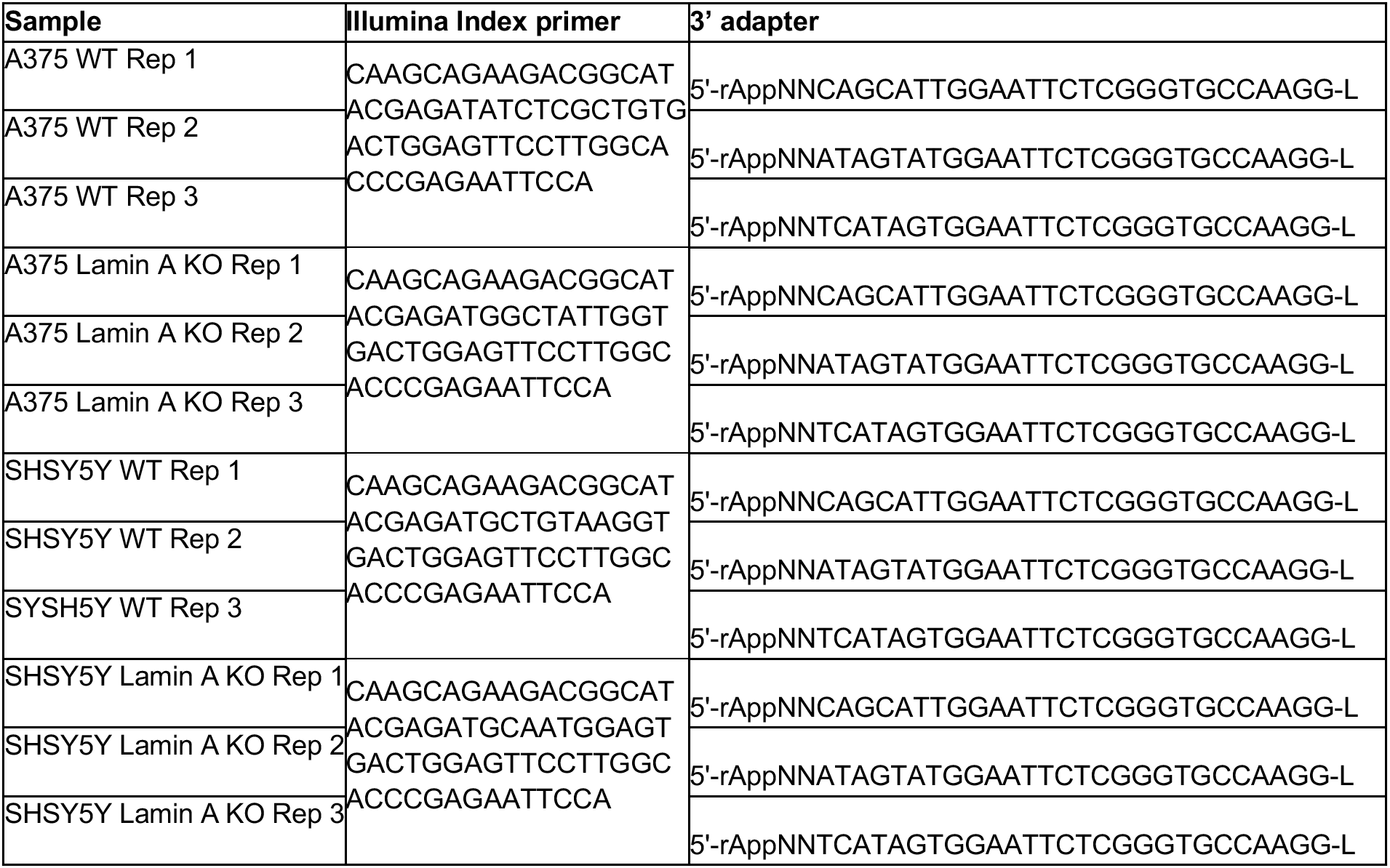

### Fluorescent PhotoActivatable Ribonucleoside-enhanced CrossLinking and ImmunoPrecipitation (fPAR-CLIP)

AGO or FAM120A fPAR-CLIP was carried out by isolating the proteins using T6B peptide as mentioned above or anti-FAM120A. Following, fPAR-CLIP library preparation, sequencing and initial data processing was performed as described in [52] with minor modifications. Briefly, to obtain protein RNA footprints, unprotected RNA was digested on beads with 1 U RNase T1 (EN0541, ThermoFisher) for 15 min at RT. Next, the beads were washed three times with RIPA buffer and three times with dephosphorylation buffer (50 mM Tris-HCl–HCl, pH 7.5, 100 mM NaCl, 10 mM MgCl_2_). After washing, the protein bound RNA was dephosphorylated with Quick CIP (M0525S, New England Biolabs) for 10 min at 37°C. Post dephosphorylation the beads were washed three times with dephosphorylation buffer and three times with PNK/ligation buffer (50 mM Tris-HCl-HCl, pH 7.5, 10 mM MgCl_2_). Following, 0.5 μM fluorescently tagged 3’ adapter (MultiplexDX) were ligated with T4 Rnl2(1–249)K227Q (M0351, New England Biolabs) overnight at 4°C and washed three times with PNK/ligation buffer. Next, RNA footprints were phosphorylated using T4 PNK (NEB, M0201S) for 30 min at 37°C and washed three times with RIPA buffer. To elute the proteins, the beads were incubated at 95°C for 5 min in 2× SDS Laemmli buffer. Next, the eluates were separated on a 4-12% SDS/PAGE gels (NW04122BOX, Invitrogen) and AGO:RNA complexes visualized on the IR 680 channel (Chemidoc MP system, Bio Rad). Subsequently, appropriate AGO:RNA bands were excised from the gel, protein digested with Proteinase K (RPROTK-RO, Sigma Aldrich) and released RNA isolated via phenol:chlorophorm phase separation. Following, 5’ adapter ligation (MultiplexDX) was performed on the purified RNA samples with 0.5 μM of the adapter and Rnl1 T4 RNA ligase (EL0021, ThermoFisher) for 1 h at 37°C. Next, the RNA was reverse transcribed using SuperScript IV Reverse Transcriptase (18090010, ThermoFisher) according to manufacturer’s instructions. The libraries were amplified in a series of PCR reactions performed using Platinum Taq DNA polymerase (10966034, ThermoFisher) and size selected with 3% Pippin Prep (CSD3010, Sage Science). Sequencing of the libraries was carried out on Illumina NovaSeq 6000 platform. For data processing Bcl2fastq (v2.20.0), Cutadapt (cutadapt 1.15 with Python 3.6.4) [54], PARpipe (https://github.com/ohlerlab/PARpipe) and Paralyzer [55] were used. The 3’ and 5’ adapter sequences and sequencing primers used in the study are listed below.

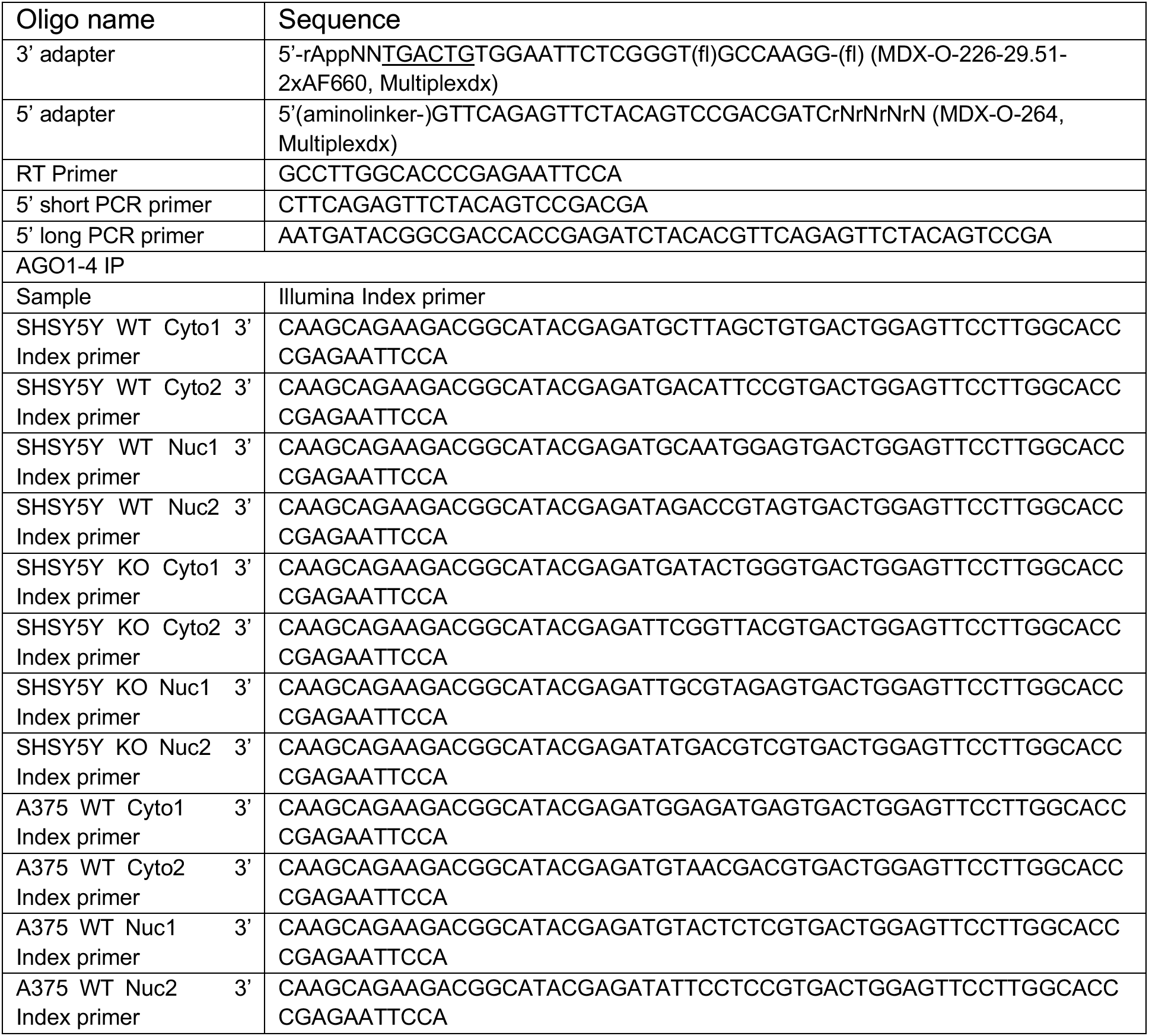
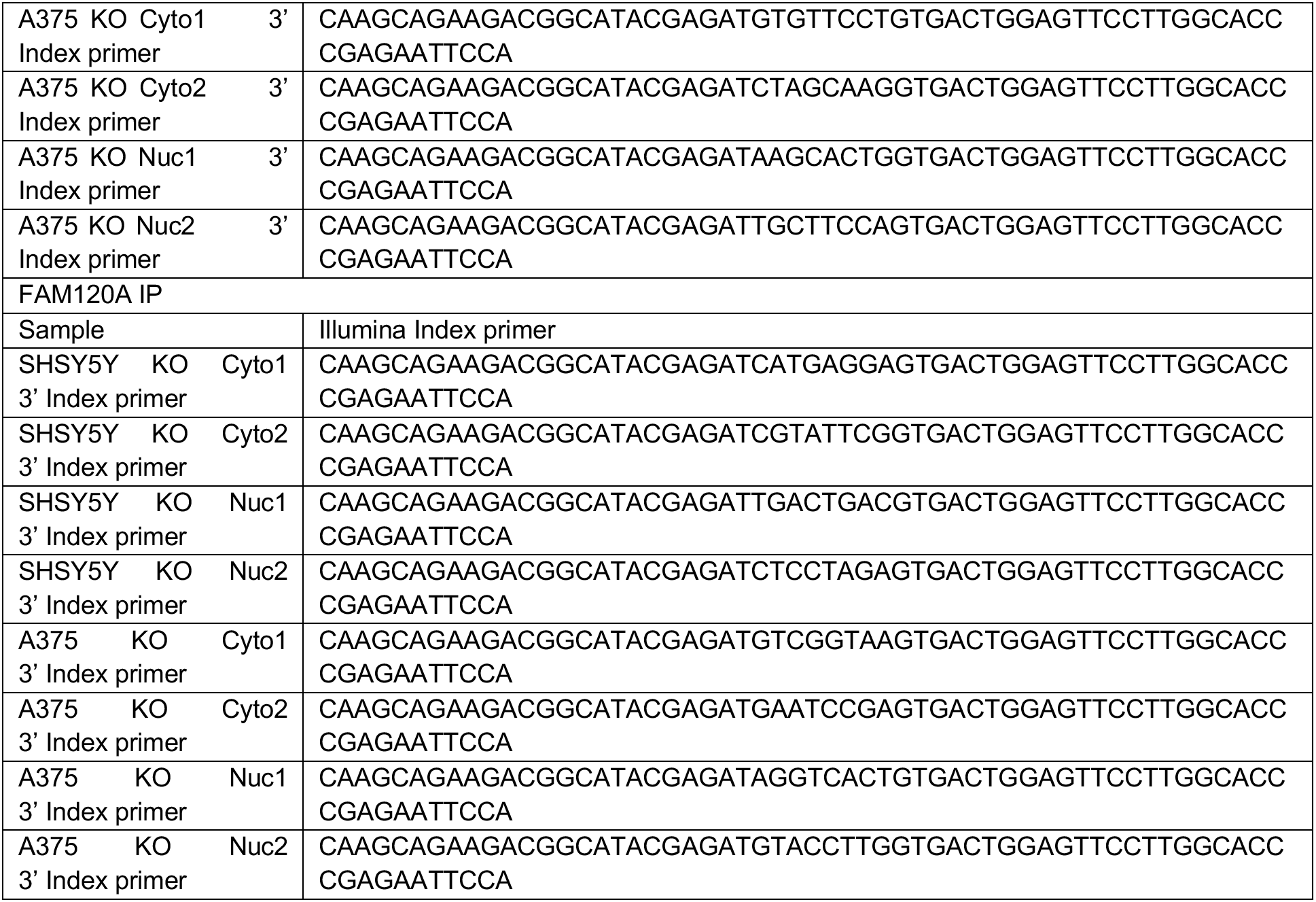

### Oxidative stress assessment

For detection of H_2_O_2_ species in cell culture media of SHSY5Y and A375 Lamin KO and WT cells, as well as SHSY5Y and A375 Lamin KO cells transfected with pCDH-CMV-hLamin_A-IRES-copGFP-EF1-puro plasmid, ROS-Glo™ H_2_O_2_ Assay (G8820, Promega) was used. Additionally, to trigger intracellular ROS production, cells were treated with 25 μM Menadione (M5625, Sigma Aldrich) for 3 h. Furthermore, ROS quenching was carried out by treating cells with 5 mM N-acetyl cysteine (NAC) (616-91-1, Merck) for 1 h.

### Cell viability evaluation

For assessment of cell viability cells were seeded in 96-well plates and 24 h after the redox potential of the cells was measured using CyQUANT™ MTT Cell Viability Assay (V13154, Invitrogen) according to manufacturers instructions.

## RESULTS

### The subcellular localization of AGO2 is affected by the nuclear lamina composition

The subcellular localization of RNA binding proteins is relevant to their function. Although, AGO2 is best studied in the cytoplasm [13–14,18, 23, 56], nuclear AGO2 has also been characterized [15–26,36,56–57] raising questions regarding nuclear RNAi and additional putative functions of AGO2 in the nucleus. To probe AGO2 nuclear localization in cancer cells, we performed biochemical fractionation experiments in twelve different human cell lines and detected fluctuating levels of AGO2 between the cytoplasmic and nuclear fractions (**Figure 1A-C**). In seven of the screened cell lines, nuclear AGO2 comprised at least 30% and up to 60% of the total AGO2 protein (**Figure 1B,C**), while the other five cell lines express seemingly negligible levels of nuclear AGO2 (**Figure 1A,C**). No pattern was observed in eirther cell origin or cancer stage in the cells expressing nuclear AGO2.

**Figure 1.**
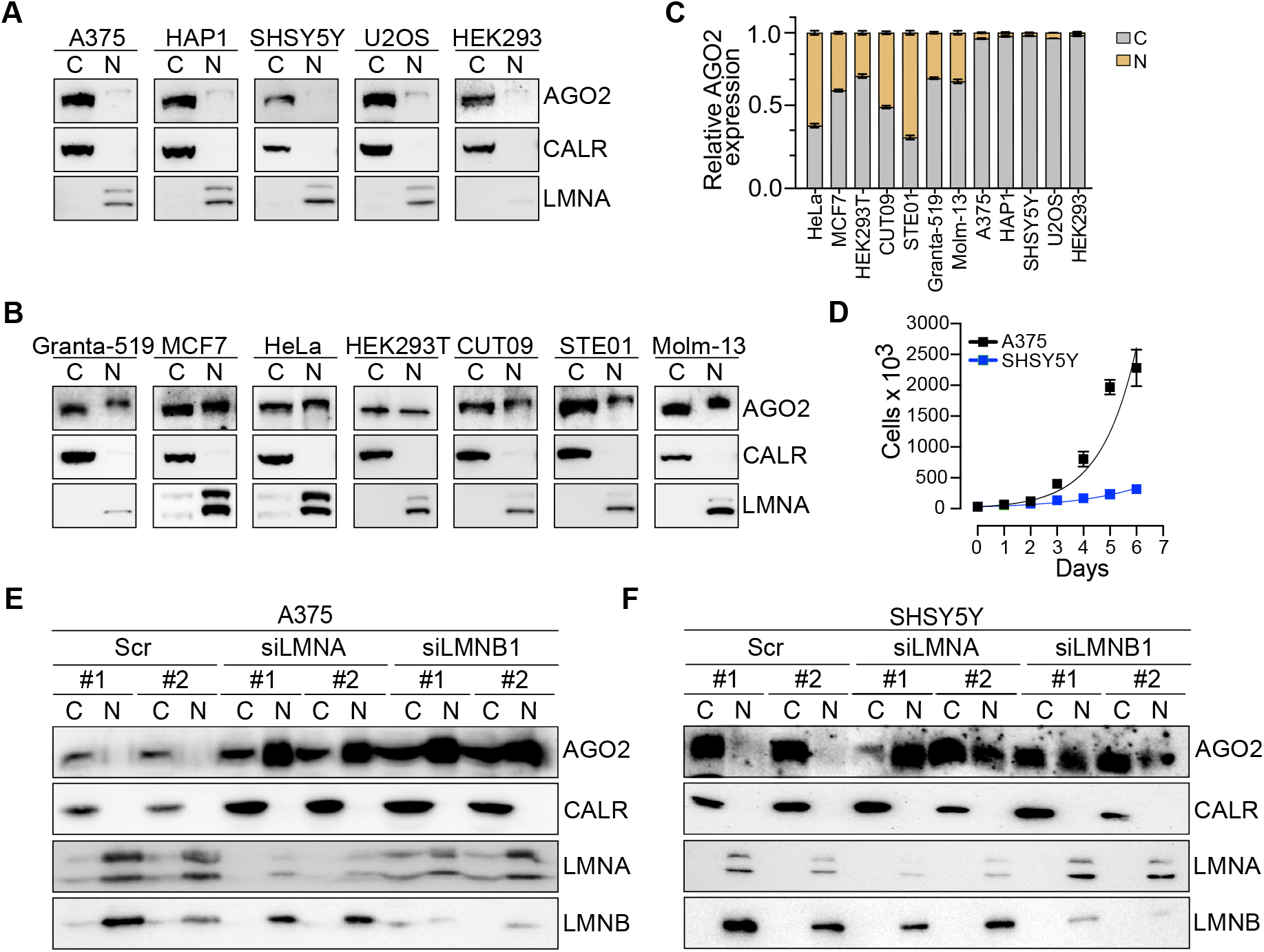
Silencing of *LMNA* or *LMNB1* expression triggers AGO2 nuclear translocation. Representative images of AGO2 immunoblots from cytoplasmic and nuclear fractions in a panel of **(A)** nuclear AGO2 negative and **(B)** nuclear AGO2 positive human cell lines. (C) Relative ratio of nuclear and cytoplasmic AGO2 fractions in the studied cell lines. **(D)** Cell proliferation assay for A375 and SHSY5Y cells. Exponential (Malthusian) growth for proliferation assays line was fitted for the data sets. Representative images of AGO2 immunoblots from cytoplasmic and nuclear fractions in cells transfected with the siRNAs for LMNA and LMNB1 in **(E)** A375 and **(F)** SHSY5Y cells. For biochemical fractionation experiments CALR served as cytoplasmic marker while LMNA and LMNB as nuclear markers. C, cytoplasmic fraction; N, nuclear fraction; CALR, Calreticulin; LMNA, Lamin A/C; Cont, cells treated with Lipofectamine RNAiMAX transfection agent; Scr, cells transfected with scrambled oligonucleotide; si*LMNA*, cells transfected with siRNAs against *LMNA*; si*LMNB1,* cells transfected with siRNAs against *LMNB1.* Graphs presented in the figure represent results calculated from three independent experiments and presented as the mean ± standard deviation.

Nucleocytoplasmic translocation of macromolecules is bridged by the NE. Proteins exceeding a molecular weight of 40 kDa are actively transported through the nuclear pore complexes (NPC), which are large multiprotein complexes, embedded in the NE [58]. We hypothesized that the NE restricts nuclear AGO2 translocation and, since the nuclear lamina is a major component of the NE, the nuclear entry of AGO2 may be promoted by reduced rigidity of the nuclear lamina meshwork. To test the possible effect of Lamin levels on AGO2 subcellular localization we opted for Lamin A/C and Lamin B1 depletion by *LMNA* and *LMNB1* targeting siRNAs (**Supplemental Figure 1A**). A375 and SHSY5Y cell lines were selected because they do not have detectable nuclear AGO2 levels (**Figure 1A,C**), and their proliferation rate and tumorigenicity differs significantly, with SHSY5Y characterized by slower proliferation (**Figure 1D**) and less tumor formation in mice [59–60]. 48 hours after siRNA transfections we performed a series of biochemical fractionation experiments in the *LMNA* and *LMNB1* siRNA-transfected cells, which revealed that siRNA-mediated reduction of Lamin A/C and Lamin B1 levels triggered AGO2 nuclear translocation in both A375 melanoma and SHSY5Y neuroblastoma cells (**Figure 1E,F**). While B type Lamins are ubiquitously expressed, Lamin A is only expressed in differentiated cells [61]. Therefore, we pursued *LMNA* depletion also in HEK293 and U2OS cells, which normally do not express nuclear AGO2, to further test the role of Lamin A/C in the cytoplasmic retention of AGO2 (**Figure 1A-C**). In both HEK293 and U2OS cell lines, the depletion of Lamin A/C resulted in the translocation of AGO2 into the nucleus (**Supplemental Figure 1B,C**).

Taken together, our results indicate that the nuclear lamina composition favors cytoplasmic localization of AGO2 in certain cancer cells, which can be reversed when Lamin levels are diminished.

### Knockout of Lamin A affects the subcellular localization of RISC components in A375 and SHSY5Y cells

The striking nuclear translocation of AGO2 in *LMNA* depleted A375, SHSY5Y, HEK293 and U2OS cells (**Figure 1E,F, Supplemental Figure 1B,C**), prompted us to further explore the correlation between *LMNA* expression and AGO2 subcellular localization. Therefore, we performed CRISPR-Cas9 to KO Lamin A in A375 and SHSY5Y, two cancer cell lines negative for nuclear AGO2, and in HeLa cervical cancer cells, which are positive for nuclear AGO2 (**Figure 1A-C**). We targeted the CRISPR-Cas9 to the constitutive exon 9 of the gene, which is common between Lamin A and Lamin C and indispensable to produce all A-type Lamins [62] (**Supplemental Figure 2A**). The Lamin A KO was verified by PCR (**Supplemental Figure 2B-D**) and by western blotting using either Lamin A/C antibody or antibody specific for Lamin A (**Figure 2A,B**). We observed complete KO of Lamin A, while, unexpectedly, low levels of Lamin C were still detectable. The functionality of the expressed form of Lamin C protein was not further evaluated. A series of immunofluorescence experiments were performed in the WT and Lamin A KO A375 and SHSY5Y cells to further confirm Lamin A protein KO and to assess the cellular morphology of the Lamin A KO cells. Consequently, we observed a lack of Lamin A expression in A375 and SHSY5Y KO cells (**Figure 2C,D**). The morphology of A375 cells did not alter significantly upon Lamin A KO, however, the loss of Lamin A resulted in complete atrophy of neurites of SHSY5Y cells (**Supplemental Figure 2E**). Furthermore, the nuclei and cytoplasm of the Lamin A KO cells were misshapen; however, the effect was more visible in the case of SHSY5Y Lamin A KO cells (**Figure 2D, Supplemental Figure 2E**). Additionally, we showed that RNA Polymerase II, Histone H3 and Lamin B1 nuclear localization, as well as cytoplasmic-localized Actin, was not affected by Lamin A KO (**Figure 2C,D, Supplemental Figure 2F,G**), which indicates that the NE at least partially functional in the Lamin A KO cells. To further exclude the occurrence of nuclear ruptures and unspecific nuclear translocation of proteins in the Lamin A KO cells, we transfected A375 and SHSY5Y WT and Lamin A KO cells with NLS-GFP constructs. NLS-GFP signal was equally confined to the nucleus in both WT and Lamin A KO cells, with minimal GFP signal observed in the cytoplasm, further pointing to the stability of the NE (**Figure 2E,F upper panel**). Furthermore, since AGO2 is associated with the Endoplasmic Reticulum (ER) and to exclude the possibility that AGO2 nuclear translocation in Lamin A KO conditions is due to ER leakage into the nucleus, we stained the ER using ER tracker (**Figure 2E,F lower panel**). Our results indicate that the integrity of the ER is not affected upon loss of Lamin A, with no observed penetrance of ER components into the nucleus (**Figure 2E,F lower panel**).

**Figure 2.**
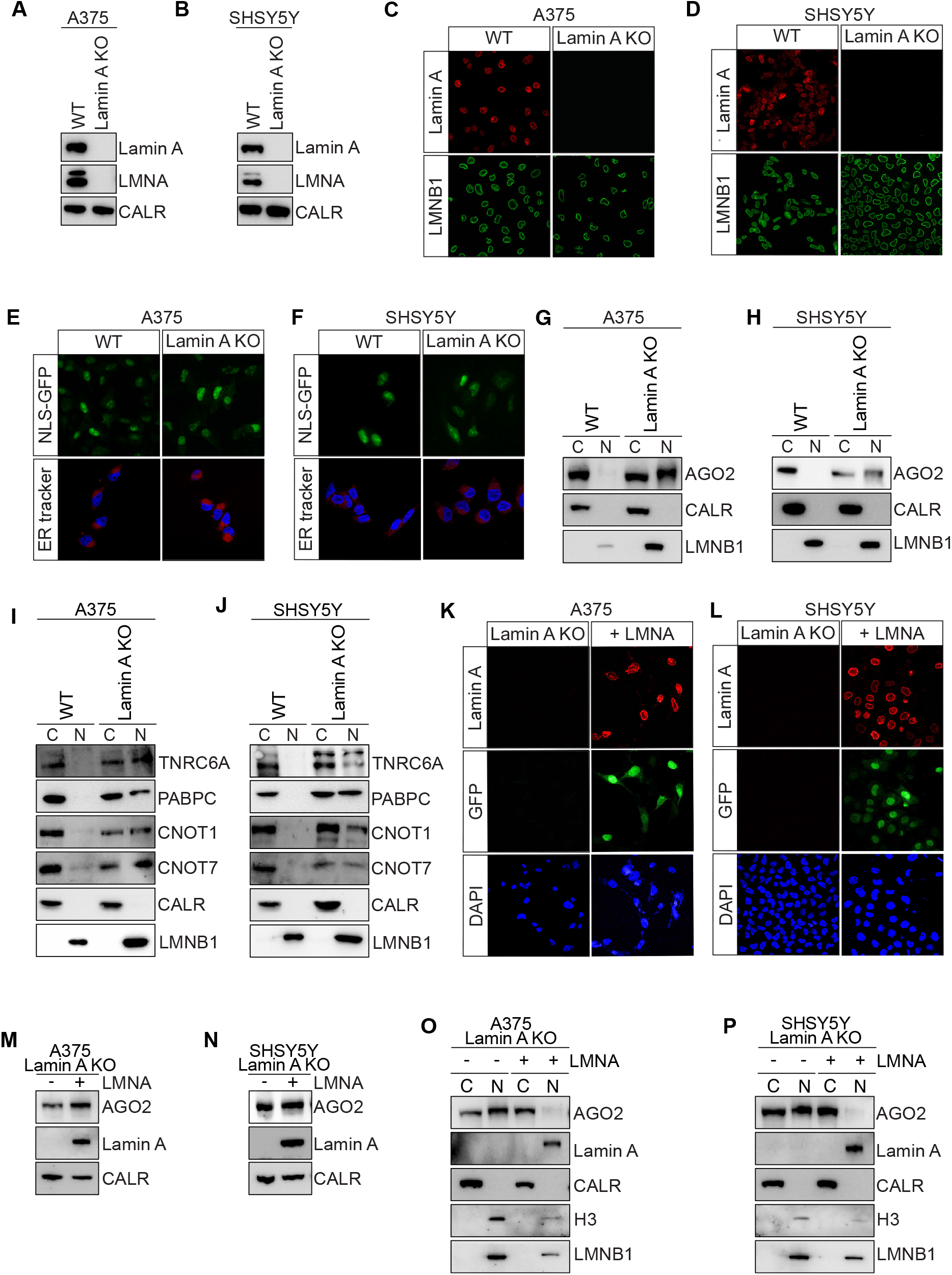
Lamin A KO causes significant nuclear translocation of RISC complex proteins. Representative images of Lamin A, Lamin A/C and Calreticulin immunoblots from **(A)** A375 and **(B)** SHSY5Y WT and Lamin A KO cells. Immunofluorescence staining of Lamin A and LMNB1 in **(C)** A375 and **(D)** SHSY5Y WT and Lamin A KO cells. Representative images of NLS-GFP fluorescent signal (upper panel) and ER tracker (lower panel) in **(E)** A375 and **(F)** SHSY5Y WT and Lamin A KO cells. Representative images of AGO2 immunoblots from cytoplasmic and nuclear fractions in **(G)** A375 and **(H)** SHSY5Y WT and Lamin A KO cells. Representative images of TNRC6, PABPC, CNOT1 and CNOT7 immunoblots from cytoplasmic and nuclear fractions in **(I)** A375 and **(J)** SHSY5Y WT and Lamin A KO cells. Immunofluorescence staining of Lamin A, GFP and DAPI in **(K)** A375 and **(L)** SHSY5Y Lamin A KO cells and Lamin A KO Lamin A/C overexpressing cells. Representative images of AGO2 and Lamin A immunoblots from **(M)** A375 and **(N)** SHSY5Y Lamin A KO and Lamin A KO Lamin A/C overexpressing cells. CALR was used as a loading control. Representative images of AGO2 and Lamin A immunoblots from cytoplasmic and nuclear fractions in Lamin A KO and Lamin A KO Lamin A/C overexpressing **(O)** A375 and **(P)** SHSY5Y cells. For biochemical fractionation experiments the endoplasmic reticulum protein Calreticulin served as cytoplasmic marker while H3 and LMNB1 served as nuclear markers. WT, wild-type cells; Lamin A KO, Lamin A knock out cells; LMNA, Lamin A/C; CALR, Calreticulin; H3, Histone H3; LMNB1, Lamin B1.

As mentioned above, siRNA-mediated silencing of Lamin A/C expression resulted in AGO2 translocation into the nucleus in several cancer cell lines (**Figure 1E,F, Supplemental Figure 1B,C**). To further explore the link between complete knockout of Lamin A and AGO2 localization, we fractionated A375 and SHSY5Y WT and Lamin A KO cells (**Figure 2G,H)**. Fractionation experiments showed predominantly cytoplasmic localization of AGO2 protein in A375 and SHSY5Y WT cells (**Figure 2G,H**). However, potent and significant localization of nuclear AGO2 was found in both A375 and SHSY5Y Lamin KO cells (**Figure 2G,H**). A similar trend was observed for the remaining AGO family proteins in both A375 and SHSY5Y Lamin KO cells (**Supplemental Figure 2H,I**). In HeLa cells that are characterized by ubiquitous distribution of AGO2 between the nucleus and the cytoplasm (**Figure 1B,C**), Lamin A KO minimally affected AGO2 localization (**Supplemental Figure 2J**). Additionally, we characterized AGO2 protein and RNA expression levels upon Lamin A KO. We observed no significant change in *AGO2* mRNA levels in A375 cells (**Supplemental Figure 2K**). In contrast, *AGO2* mRNA levels was significantly increased in SHSY5Y Lamin A KO cells (**Figure 2L**). This trend was also reflected in AGO2 protein levels, where there was significant increase in AGO2 protein level in SHSY5Y Lamin A KO cells, but not in A375 Lamin A KO cell (**Supplemental Figure 2M,N**).

AGO2 is central for RNAi processes, therefore we next wondered if key components of the RNA induced silencing complex (RISC) follow the AGO2 trend of nuclear translocation upon loss of Lamin A. Therefore, we fractionated A375 and SHSY5Y WT and Lamin A KO cells and probed for the subcellular localization of TNRC6A, PABPC and members of the CCR4-NOT deadenylation complex (CNOT1 and CNOT7). All tested RNAi factors co-translocated into the nucleus with AGO2 upon loss of Lamin A (**Figure 2I,J**).

Finally, to test if rescuing Lamin A/C expression is sufficient to restore the WT cellular phenotype of A375 and SHSY5Y cells, where no nuclear AGO2 is detected, we performed *LMNA* overexpression experiments in the Lamin A KO cells. Localization of ectopically expressed Lamin A/C at the nuclear envelope was confirmed by immunofluorescent staining using anti-Lamin A antibodies, and transfection efficiency was assessed by GFP expression (**Figure 2K,L**). Moreover, the levels of Lamin A ectopic expression were confirmed by western blot (**Figure 2M,N**). Biochemical fractionation experiments revealed a significant decrease in nuclear AGO2 content in *LMNA* overexpressing cells in both A375 and SHSY5Y Lamin A KO cells, resulting in a rescue of WT phenotype, where AGO2 has a predominant cytoplasmic distribution (**Figure 2O,P**). This indicates that nuclear import of AGO2 is directly linked with diminished levels of Lamin A.

### Loss of Lamin A increased cell growth and oncogenic miRNAs expression in SHSY5Y but less in A375 cells

We further proceeded to characterize the phenotypes of Lamin A KO cells. First, we evaluated cell proliferation and found that while A375 cell growth rate changed minimally in response to loss of Lamin A (**Figure 3A**), proliferation of SHSY5Y cells were significantly increased (**Figure 3B**), reaching growth rates comparable to A375 cells. In line with this type of investigation, we also measured cell viability (**Supplemental Figure 3A,B**) and cell cycle rates (**Supplemental Figure 3C,D**) in WT vs Lamin A KO cells. Cell viability was significantly increased after Lamin A KO (**Supplemental Figure 3A,B)**, with enrichment in G2/M phase of the cell cycle (**Supplemental Figure 3C,D**).

**Figure 3.**
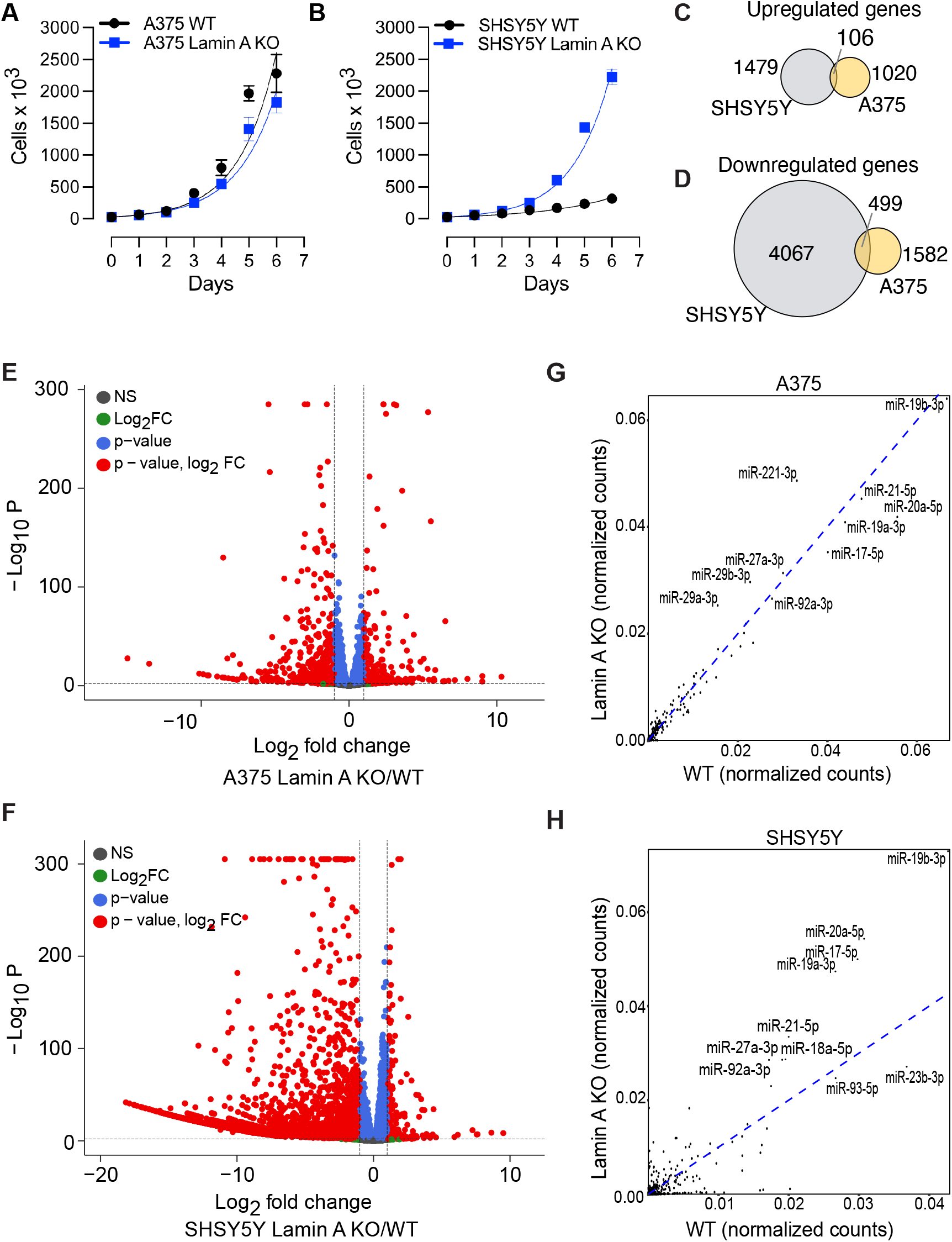
Lamin A KO triggers changes in gene expression and miRNA profile in SHSY5Y and A375 cells. Cell proliferation assay for growing Lamin A KO and WT **(A)** A375 and **(C)** SHSY5Y cells. Exponential (Malthusian) growth line for proliferation assays was fitted for the data sets. Results were calculated from four independent experiments performed in technical duplicates and presented as the mean ± standard deviation. Venn diagram of the overlap of genes **(C)** upregulated and **(D)** downregulated in SHSY5Y and A375 cells upon Lamin A KO. Volcano plot depicting the differential gene expression between WT and Lamin A KO transcripts in **(E)** A375 and **(F)** SHSY5Y cells. miRNA profile bound to AGO1-4 in WT and Lamin A KO cells in **(G)** A375 and **(H)** SHSY5Y. WT, wild-type cells; Lamin A KO, Lamin A knock out cells.

To assess the changes in gene expression between A375 and SHSY5Y Lamin A KO cells compared to their WT counterparts, we performed RNA sequencing experiments. Principal Component Analysis (PCA) revealed significant differential gene expression changes between the Lamin A KO and WT cells (**Supplemental Figure 3E,F**). In A375 we observed 1020 upregulated genes and 1582 downregulated genes (**Figure 3C,D**). In SHSY5Y we found 1479 upregulated genes and 4067 downregulated genes (**Figure 3C,D**). The differences in gene expression between Lamin A KO and WT cells, were depicted in a volcano plot (**Figure 3E,F**, **Table 1**) and a GO pathway analysis was performed for the top 100 differentially expressed genes (**Supplemental Figure 3G**). Our overall results reveal a global downregulation of gene expression, which is exaggerated in SHSY5Y cancer cells.

miRNA regulation has been shown to be modified in cancers where individual miRNA species can either act as tumor suppressor or oncogenic factors [10,27]. To understand how miRNAs respond to loss of Lamin A, we profiled the miRNA populations bound to AGO1-4 in WT and Lamin A KO cells. AGO1-4 was immunoprecipitated using the T6B peptide and the bound small RNAs were isolated and deep sequenced. The miRNA profile in A375 were similar in WT and Lamin A KO cells (**Figure 3G**; **Table 2**), however, we found that miRNAs were strikingly differentially expressed in SHSY5Y cells upon Lamin A KO (**Figure 3H**; **Table 2**). This result is in accordance with the observed effects of Lamin A KO on cell growth in the two cell lines. Of note, in SHSY5Y cells, we observed potent upregulation of several miRNAs encodes by the miR-17/92 cluster, such as miR-19b, miR-19a, miRNA-18a, miRNA-92a miR-17 and miR-20a, upon Lamin A KO (**Figure 3H**). The increase in miRNA expression in Lamin A KO conditions, was also accompanied by increased AGO2 protein levels in SHSY5Y cells (**Supplemental Figure 2N**).

Our high-throughput transcriptomics experiments collectively suggest different mechanisms of gene regulation in A375 and SHSY5Y cells to cope for the loss of Lamin A, however, both mechanisms lead to a potent accumulation of AGO2 in the nucleus. The loss of Lamin A has a more profound phenotypic effect in SHSY5Y cells, which are cancer cells characterized with less growth potential (**Figure 1D**). In A375, the loss of Lamin A effect is less noticeable, indicating that growth rates cannot be further affected by the loss of Lamin A. The nuclear translocation of AGO2 may therefore be regulated by different cellular mechanisms depending on the cancer cell type.

### RNAi activity is significantly diminished in Lamin A KO cells

Having determined that there is substantial translocation of AGO2 into the nucleus in cancer cells when Lamin A levels are diminished, we aimed at understanding the putative functions of AGO2 in the nucleus. AGO2, guided by miRNAs, mediates gene regulation in the cytoplasm by targeting mRNAs predominantly within coding regions and 3’ untranslated (3’UTR) regions [12]. Previous studies provided evidence that natural cell conditions, such as viral infection [63–64] or early differentiation [18], comprise functional AGO2 nuclear regulation and that AGO2 preferentially binds 3’UTRs and introns within pre-mRNAs [18,64]. To gain detailed information on AGO-dependent regulation on gene expression in Lamin A KO cells we performed fPAR-CLIP in WT and Lamin A KO A375 and SHSY5Y cells. First, cells were fractionated and the fractions, as well as efficiency of immunoprecipitation was assessed (**Figure 4A, Supplemental Figure 4A**). The crosslinked and purified ribonucleoprotein (RNP) complexes were visualized by SDS-PAGE fluorescence imaging, which revealed a band at ∼ 130kDa corresponding to AGO1-4 RNP (**Figure 4B, Supplemental Figure 4B**).

**Figure 4.**
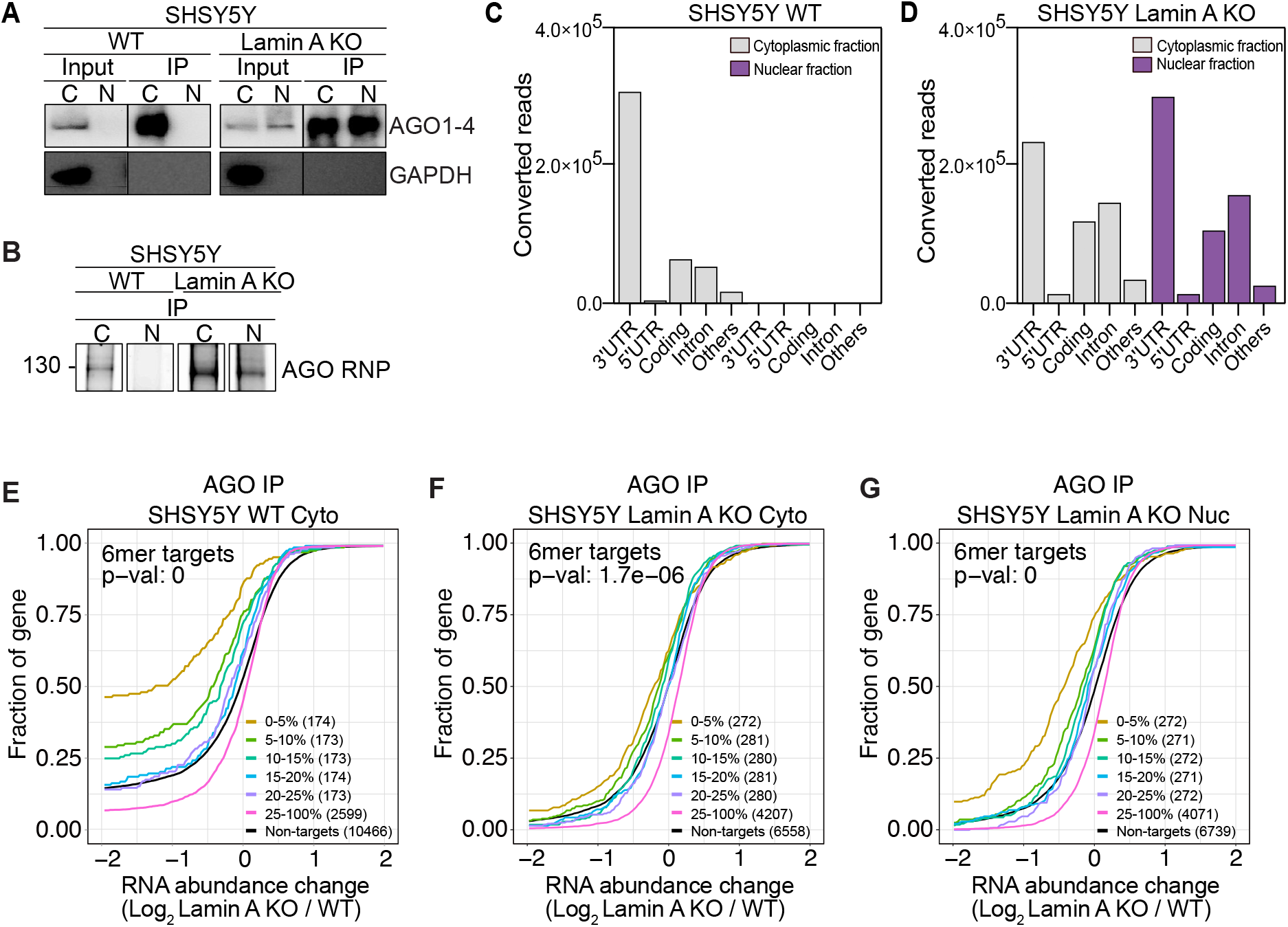
Nuclear RNAi is diminished upon Lamin A KO. (A) Representative images of AGO2 and GAPDH immunoblots from SHSY5Y WT and Lamin A KO cells after immunoprecipitation of AGO1-4 using the T6B peptide. (**B**) Fluorescence SDS-PAGE of 3’ labeled AGO1-4 RNP in WT and Lamin A KO SHSY5Y cells. Distribution of AGO fPAR-CLIP sequence reads across target RNAs in SHSY5Y **(C)** WT cytoplasmic and nuclear lysate and **(D)** Lamin A KO cytoplasmic and nuclear lysate. Cumulative distribution of abundance changes in RNA in SHSY5Y **(E)** WT cytoplasmic fraction **(F)** Lamin A KO cytoplasmic fraction and **(G)** Lamin A KO nuclear fraction. For cumulative distribution assays the targets were ranked by number of binding sites from top 5-20% and 25-100% and compared to non-targets. WT, wild-type cells, Lamin A KO, Lamin A knock out cells; C, cytoplasmic fraction; N, nuclear fraction; IP, immunoprecipitation; fPAR-CLIP, fluorescent photoactivatable ribonucleoside-enhanced crosslinking and immunoprecipitation; RNP, ribonucleoprotein complex.

In the cytoplasmic fraction of WT cells, AGO occupied predominantly the 3’UTR region of mRNAs (**Figure 4C, Supplemental Figure 4C, Table 3**). As expected, due to the cytoplasmic localization of AGO2 in WT A375 and SHSY5Y cells, we observed minimal transcript occupancy of AGO in the nuclear fraction of both SHSY5Y (**Figure 4C**, **Table 3**) and A375 cells (**Supplemental Figure 4C, Table 3**). On the other hand, upon Lamin A KO we observed less specific occupancy of the AGOs, which more evenly distributed between 3’UTR, introns, and coding regions (**Figure 4D, Supplemental Figure 4D, Table 3**), reflecting the shift in AGO subcellular localization. However, to our surprise, nuclear and cytoplasmic AGOs were not distinct in their target RNAs occupancy, where also cytoplasmic AGO proteins targeted introns in the Lamin A KO cells (**Figure 4D, Supplemental Figure 4D**).

To estimate the RNAi potential of nuclear AGOs we integrated the RNAseq and miRNAseq results from Lamin A KO and WT cells with the AGO fPAR-CLIP data. Firstly, we ranked genes by their AGO occupancy in WT cells and used cumulative distribution analysis to test the effect of Lamin A KO on their expression level (**Figure 4E, Supplemental Figure 4E**). Only AGO targets containing a 6mer seed sequence for the top 200 highest expressed miRNAs (accounting for <95% of miRNA expression) were considered. The analysis revealed that specifically AGO targets in cytoplasmic fraction of WT SHSY5Y cells are increasingly downregulated in response to Lamin A KO, and directly proportional to the binding strength of AGO (**Figure 4E**). Unlike AGO targets in WT cells, the top targets acquired by AGO in Lamin A KO cells, from cytoplasmic and nuclear fractions, were generally less downregulated in response to Lamin A KO, as compared to WT cells (**Figure 4F,G**). Contrary to the potent RNAi effect observed in WT SHSY5Y cells, in A375, the AGO mediated RNAi was negligible in both WT and Lamin A KO cells (**Supplemental Figure 4E-G**), indicating that miRNA pathways are less active in the highly proliferative A375 cells.

Taken together, our analyses of AGO-dependent gene regulation in Lamin A KO vs WT cells revealed that the pool of AGOs which translocate into the nucleus does not predominantly and effectively engage in gene regulation. This was surprising as we observed that RISC components, such as TNRC6, needed for RNAi, translocated to the nucleus in Lamin A KO cells (**Figure 2I,J**). Therefore, there may be other mechanisms of gene regulation at play in the Lamin A KO cells.

### Loss of Lamin A promotes nuclear AGO2 interaction with FAM120A, nucleoporins and chromatin regulators

To gain more insights into the nuclear functions of AGO1-4, we investigated the interactome of nuclear AGO proteins by mass spectrometry using the T6B peptide, which unbiasedly binds all four AGO proteins [41]. We identified 184 AGO interactors enriched in the A375 Lamin A KO nuclei, compared to WT nuclei, which do not express nuclear AGO and therefore are treated as a negative control (**Figure 5A, Supplemental Table 4**). Respectively, in the SHSY5Y Lamin A KO nuclei, compared to WT nuclei, we identified 141 AGO interacting partners (**Figure 5B, Supplemental Table 4**). Gene ontology (GO) analysis of the nuclear interacting proteins revealed that they are predominantly involved in various aspects of RNA processing (**Supplemental Figure 5A,B**).

**Figure 5.**
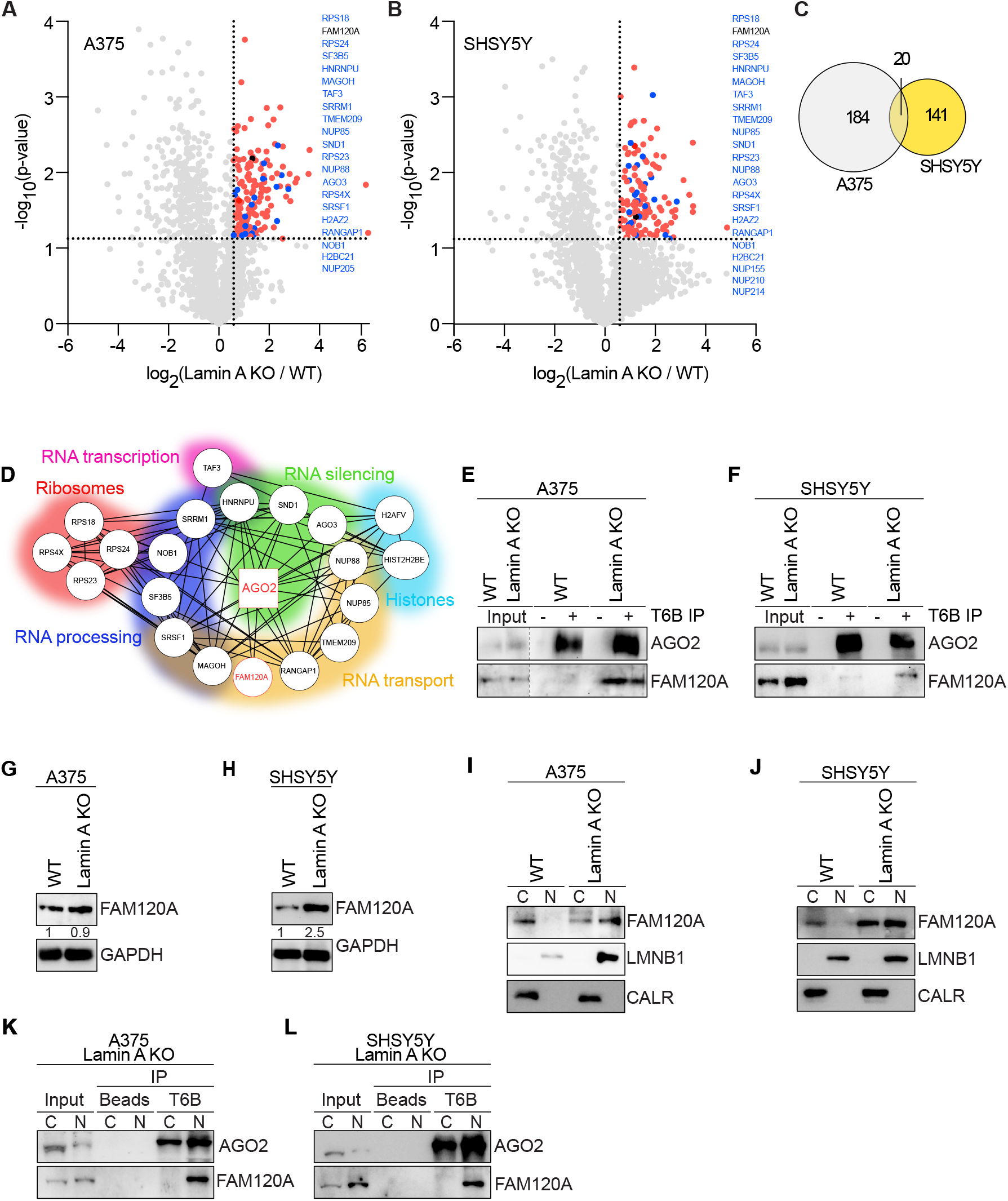
AGO2 interacts with proteins involved in RNA processing, transport in the nucleus of Lamin A KO cells and FAM120A. Volcano plots of mass spectrometry hits in Lamin A KO vs WT **(A)** A375 and **(B)** SHSY5Y nuclear lysates. Proteins significantly upregulated in Lamin A KO nucleus are indicated in red, blue and black. **C)** Venn diagram of significantly upregulated mass spec hits in the nucleus of Lamin A KO vs WT A375 (indicated in grey) and SHSY5Y (indicated in yellow) cells. **(D)** String analysis of significantly upregulated mass spec hits in the nucleus of Lamin A KO vs WT overlap between A375 and SHSY5Y cells. Representative images of AGO2 and FAM120A immunoblots from co-immunoprecipitation assay in **(E)** A375 and **(F)** SHSY5Y cells. Representative images of FAM120A immunoblots from whole cell lysates in **(G)** A375 and **(H)** SHSY5Y cells. Representative images of FAM120A immunoblots after biochemical fractionation into cytoplasmic and nuclear fractions in **(I)** A375 and **(J)** SHSY5Y cells. Calreticulin was used as a cytoplasmic marker while Lamin B1 served as a nuclear marker. Representative images of AGO2 and FAM120A immunoblots from AGO1-4 immunoprecipitation assay in cytoplasmic and nuclear lysates from **(K)** A375 and **(L)** SHSY5Y WT and Lamin A KO cells. WT, wild-type cells; Lamin A KO, Lamin A knock out cells; C, cytoplasmic fraction; N, nuclear fraction; CALR, calreticulin; LMNB1, Lamin B1; T6B IP, immunoprecipitation using T6B peptide.

Further examination of the enriched interacting partners of nuclear AGO revealed that there are 20 protein interactors which overlapped between A375 and SHSY5Y cells (**Figure 5C**). Functional annotation analysis of this subset of interacting partners further confirmed that they are involved in regulation of various aspects of RNA processing (**Figure 5D**). The list also included nuclear pore complex (NPC) components Nup85, Nup88 and four DNA associated regulators, including Histone H2B, H2A.V, and the transcription initiation factor TFIID. Interestingly we also identified the Constitutive coactivator of PPAR-gamma-like protein 1, FAM120A, as an interacting partner of AGO in the nucleus (**Figure 5D, Supplemental Table 4**). FAM120A is an RNA binding protein involved in oxidative stress response [65]. FAM120A has previously been suggested to interact with AGO2 and act as a competitor for RNA binding [66–67]. We further validated this hit by performing co-immunoprecipitation assay between AGO2 and FAM120A from whole cell lysates (**Figure 5E,F**). In both A375 and SHSY5Y cells, AGO2:FAM120A interaction occurred in Lamin A KO cells, but not in WT cells (**Figure 5E,F**). The total levels of FAM120A was significantly increased upon Lamin A KO in SHSY5Y cells, but not in A375 cells (**Figure 5G,H**). Moreover, subcellular fractions of Lamin A KO and WT A375 and SHSY5Y cells revealed that FAM120A subcellular localization follows the trend previously observed for AGO2 (**Figure 2G,H)**, where in WT cells it is exclusively cytoplasmic, while in Lamin A KO cells it is ubiquitously localized (**Figure 5I,J**). Nevertheless, the interaction between AGO proteins and FAM120A in the Lamin A KO cells specifically occurs in the nuclear fraction (**Figure 5K,L**)

The shift towards nuclear localization together with the identification of AGOs in complex with nucleoporins prompted us to evaluate the proximitome of the NPC using proximity labeling procedures [68]. Proximity labeling and subsequent purification allows for stringent purification of proteins from specific subcellular localizations, without biochemically fractionating cells. Therefore, we established SENP2-APEX2 expressing A375 WT and Lamin KO cells. The correct localization of SENP2-APEX2 was visualized to the NPC (**Supplemental Figure 5C**). Further, we verified that V5-tagged SENP2-APEX2 protein was expressed and active (**Supplemental Figure 5D**). By performing co-immunoprecipitation assay between biotin and AGO2 we observed an enriched proximal localization of AGO2 to the NPC in the Lamin A KO cells, compared to WT cells (**Supplemental Figure 5E**).

Additionally, we speculated that AGO2 may interact with Lamin A in Hela cells, which are normally positive for nuclear AGO2 (**Figure 1B,C**). Indeed, we found that AGO2 is in complex with Lamin A (**Supplemental Figure 5F**). Moreover, we analyzed Lamin A interactome by mass spectrometry in A375 WT and Lamin A KO cells to find possible indications of pathways that may be altered upon loss of Lamin (**Supplemental Table 4**). The GO analysis of Lamin A interacting partners revealed involvement in DNA replication and cell division processes (**Supplemental Figure 5G**), which is in line with the observed phenotype of Lamin A KO cells.

Decrease in Lamin A levels is evident in cancer progression and cellular migration, contributing to cancer aggressiveness [69]. In this condition, the sum of our findings indicates that a portion of AGO2 redistributes to the nucleus, where it interacts with RNA processing factors, possibly facilitating the cancer phenotype.

### FAM120A competes for AGO2 target binding and diminishes RNAi in Lamin A KO cells

A study by Kelly, Suzuki and colleagues in 2019 identified FAM120A as a novel interactor of Ago2 in mouse embryonic stem cells (mESC) [67]. They found that FAM120A occupies predominantly 3’ UTRs of mRNAs and interestingly, they revealed that a third of all Ago2 targets in mESC were also co-bound by FAM120A [67]. These Ago2:FAM120A shared targets were stabilized and not subjected to Ago2 mediated target suppression [67]. In Lamin A KO cells we observed potent nuclear influx of AGO2, yet to our surprise, the effect of RNAi was significantly diminished as compared to WT cells. Therefore, we next aimed to investigate if FAM120A is involved in compromising RNAi upon Lamin A KO.

Since we observed a profound increase in FAM120A protein levels in SHSY5Y (**Figure 5H**) and nuclear translocation in both SHSY5Y and A375 (**Figure 5I,J**) we next investigated if the loss of Lamin A causes increase in oxidative stress response, a condition which has previously been shown to activate FAM120 [65]. We measured the levels of hydrogen peroxide (H_2_O_2_) in WT vs Lamin A KO cells at steady state levels but also in response to Menadione, an agent which induces oxidative stress (**Supplemental Figure 6A,B**). The specificity of Menadione was further validated by using N-acetyl L cysteine (NAC), a specific inhibitor of oxidative stress. We observed an increase in H_2_O_2_ levels in Lamin A KO cells, which was further highlighted upon the addition of Menadione (**Figure 6A,B**), indicating that the Lamin A KO cells are more sensitive to oxidative stress response. To confirm that the sensitivity of oxidative stress is not an artifact of CRISPR gene editing, we also measured the oxidative stress in Lamin A KO cells where Lamin A was rescued by overexpressing Lamin A transiently. Our data shows that there is increased oxidative stress in Lamin A KO cells, compared to rescue cells (**Supplemental Figure 6C,D**).

**Figure 6.**
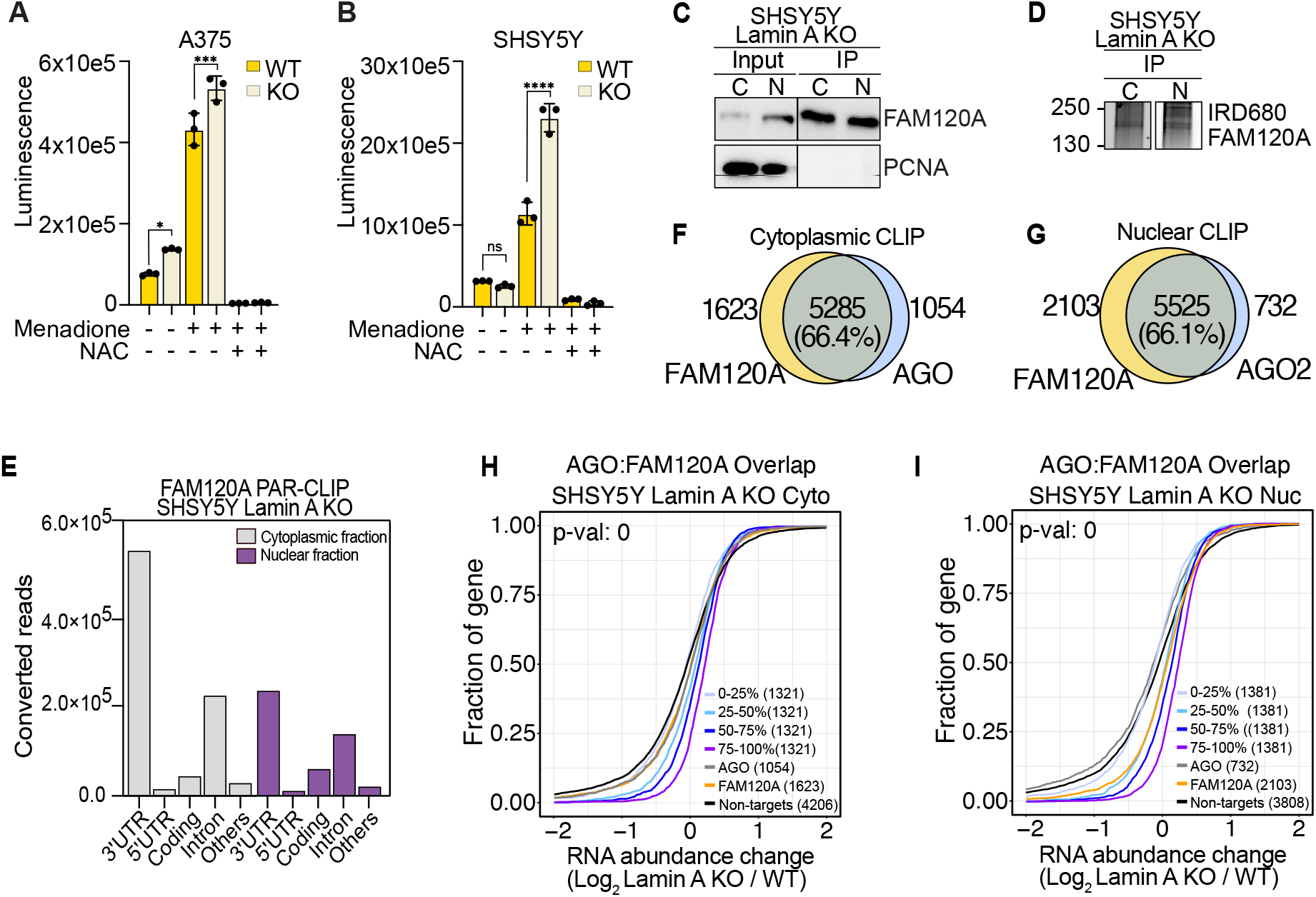
FAM120A activation upon oxidative stress leads to competitive binding of AGO targets. Assessment of H_2_O_2_ species in the media of Lamin A KO and WT **(A)** A375 and **(B)** SHSY5Y cells after treatment with ROS-inducer (Menadione) or ROS quencher (NAC). Graph present average of three independent experiments, performed in technical triplicates, mean ± standard deviation. **(C)** Representative images of FAM102A and PCNA immunoblots from SHSY5Y WT and Lamin A KO cells before and after immunoprecipitation of FAM120A. **(D)** Fluorescence SDS-PAGE of 3’ labeled FAM120A RNP in WT and Lamin A KO SHSY5Y cells. **(E)** Distribution of FAM120A fPAR-CLIP sequence reads across target RNAs in SHSY5Y Lamin A KO cytoplasmic and nuclear lysate. Overlap between AGO and FAM120A fPAR-CLIP targets in SHSY5Y cells from **(F)** cytoplasmic fraction and **(G)** nuclear fraction. Cumulative distribution of abundance changes in RNA of AGO:FAM120A overlapping targets in SHSY5Y Lamin A KO **(H)** cytoplasmic fraction and **(I)** nuclear fraction. For cumulative distribution assays the targets were ranked by 25 percentile and compared to non-targets. WT, wild-type cells, Lamin A KO, Lamin A knock out cells; C, cytoplasmic fraction; N, nuclear fraction; IP, immunoprecipitation; fPAR-CLIP, fluorescent photoactivatable ribonucleoside-enhanced crosslinking and immunoprecipitation; RNP, ribonucleoprotein complex.***P<0.001, **** P<0.0001.

Next, we performed FAM120A fPAR-CLIP from cytoplasmic and nuclear fraction of Lamin A KO cells in both A375 and SHSY5Y cells. First, the fractions, as well as efficiency of immunoprecipitation was assessed (**Figure 6C, Supplemental Figure 6E**). The crosslinked and purified ribonucleoprotein (RNP) complexes were visualized by SDS-PAGE fluorescence imaging, which revealed a band at ∼ 160kDa corresponding to FAM120A RNP (**Figure 6D, Supplemental Figure 6F**). FAM120A, similarly to AGO, occupies predominantly 3’UTR of target RNAs in the cytoplasm but revealed a more evenly distributed gene occupancy in the nucleus (**Figure 6E, Supplemental Figure 6G, Table 3**). We estimated the overlap of binding sites between AGO and FAM120A fPAR-CLIPs and observed between 60-70% overlap in both the cytoplasmic and nuclear fraction of SHSY5Y and A375 cells (**Figure 6F,G, Supplemental Figure 6H,I**). Finally, we evaluated the stability of the gene targets bound only to AGO, only to FAM120A or bound by both. The common targets were binned according to AGO binding preference and in SHSY5Y cells we found that co-bound genes were increasingly stabilized, directly proportionate to the binding strength of AGO (**Figure 6H,I**). However, in A375, the effect of AGO:FAM120A competition could not be established, even though more than 60% of all genes were co-bound (**Supplemental Figure 6J,K**), the data again reiterating the overall decreased RNAi potential in A375 cells (**Supplemental Figure 4E-G**).

Taken together, our data indicates that once AGO2 and FAM120A translocates to the nucleus in response to loss of Lamin A, FAM120A co-binds AGO2 target transcripts and stabilizes them, effectively shutting down RNAi.

## DISCUSSION

Since the discovery of RNAi, the prevalent view has been that RNAi processes are executed in the cytoplasm. RNAi-mediated silencing of nuclear transcripts was described as significantly harder to achieve as compared to cytoplasmic RNAs [13–14]. Moreover, immunofluorescent-based imaging assays repeatedly showed cytoplasmic AGO2 localization, with enrichment in P-bodies and endoplasmic reticulum [70–71]. However, improved biochemical fractionation techniques and imaging assays convincingly showed the nuclear presence of both AGO proteins, small RNAs, and other RNAi factors [15,18,20,22,36–37,72]. AGO2:miRNA complexes actively shuttle to the nucleus [15,20,22,64] and the nuclear localization of AGO2 has been shown in several cell lines, including HeLa and MCF7 used in our study (**Figure 1B,C**) [15,20,22–23,57,73] and embryonic stem cells [18]. However, the nuclear localization of AGO2 is highly dynamic and in cancer cells, such as U2OS and HAP1, AGO2 localization is predominantly cytoplasmic (**Figure 1A,C**) [18,23,56], suggesting that the fluctuations of nuclear AGO2 are tightly regulated. Yet, the mechanisms regulating the nuclear localization of AGO2, or its mode of entry, remains largely unknown. Furthermore, little connection has been made between the subcellular localization of AGO and its effect on cancer cell growth. Recently, a report showed a shift from exclusively cytoplasmic to ubiquitous localization of AGO2 in normal vs. malignant colon tissues [20], suggesting possible tumor-promoting functions of nuclear localized AGO2.

Nucleocytoplasmic translocation requires passing through the NE. The NE composition was found to be highly dynamic in cellular differentiation. While B-type Lamins are steadily expressed, Lamin A/C is expressed at very low levels in stem cells but increases during differentiation [61,73–74]. High levels of nuclear AGO2 have been shown in mouse and human stem cells [18], pointing to a possible correlation between nuclear AGO2 localization and Lamin A/C expression. However, the involvement of the nuclear lamina in the regulation of human AGO protein localization and function has not been demonstrated before. In this study, we found a relationship between Lamin A levels and AGO2 subcellular localization in cancer cells. Different experimental approaches pointed to a potent translocation of AGO2 into the nucleus in SHSY5Y neuroblastoma, A375 melanoma cells, HEK293 kidney cells, and U2OS bone cancer cells upon silencing of LMNA expression (**Figure 1E,F**, **Figure 2G,H, Supplemental Figure 1B,C**). The nuclear translocation was completely reversed when Lamin A expression was transiently rescued (**Figure 2O,P**). Thus, nuclear AGO2 expression, in the studied cell lines, seems to be directly dependent on low A-type Lamins levels.

Moreover, Lamins are speculated to play a role in cancer progression and to be altered in response to it [9,75]. LMNA is either mutated or significantly decreased in breast cancer, ovarian cancer, primary gastric carcinoma, and colon cancer [76–77]. Consistent with reports pointing to tumor-promoting functions of decreased A-type Lamins, our experiments showed increased viability of A375 and SHSY5Y cancer cells in response to Lamin A KO (**Supplemental Figure 3A,B**). Furthermore, we observed a significant increase in cell proliferation of SHSY5Y cells (**Figure 3B**), while little change in cell proliferation was observed in A375 cells, indicating that A375 cells have reached a plateau in proliferation, which cannot further be affected by decreased Lamin A levels (**Figure 3A**).

The activity of AGO proteins in the nucleus of cancer cells remain to be fully defined. In the nucleus AGO proteins have been implicated in the regulation of RNA stability, but also in chromatin remodeling, transcriptional gene regulation, alternative splicing, DNA repair, transposon silencing, apoptosis inhibition, and regulation of telomerase activity [15–20,25–26]. Of note, several studies convincingly showed efficient RNAi in the cell nuclei of embryonic stem cells, as well as human and murine cancer cells, with various methods, including CLIP and RIP techniques, proteomic and transcriptomic studies, and immunofluorescence [18,22,78–79]. Despite these findings, other report suggested inhibition of the canonical RNAi pathway in the nucleus [20]. To understand nuclear RNAi activity upon Lamin A KO, we performed fPAR-CLIP (**Figure 4, Supplemental Figure 4**). In the case of SHSY5Y cells, we observed significant RNAi impairment in Lamin A KO cells as compared to control cells (**Figure 4E-G**). Interestingly, the inhibition was found both in the cytoplasm and nucleus, suggesting an overall RNAi suppression upon Lamin A knockout, not directly connected to the cellular localization of AGO proteins (**Figure 4F,G**). Since SHSY5Y Lamin A KO cells were characterized by a more tumorigenic phenotype (**Supplemental Figure 2E, Figure 3B, Supplemental Figure 3B,D**), the observed RNAi inhibition might mimic the downregulated miRNA pathways found in aggressive cancer tissues and cell lines [27–31], rather than reflect nuclear AGO activity. This is suggestive of the RNAi effect we observed in A375, which was minimal and indicative of the fact that the miRNA pathway is not highly regulated in A375 melanoma cancer cells, as in SHSY5Y neuroblastoma cells.

The execution of RNAi dependent gene regulation is directly linked to AGO proteins and their miRNA co-factors [11–12]. Although cancer cells are characterized by the global downregulation of miRNA expression and inhibition of miRNA biogenesis, several miRNA species have been described as oncogenes and are upregulated in malignant tissues [10]. In an effort to understand the RNAi deregulation we observed upon Lamin A loss in more detail, we sequenced the miRNA species in WT and Lamin A KO cells. Similar to fPAR-CLIP, we observed a high degree of overlap in the miRNA profiles in A375 Lamin A KO and WT cells (**Figure 3G**). This once again stressed the lack of a strong phenotypic and genotypic effect of Lamin A loss on highly proliferative cancer cells. On the contrary, in the case of SHSY5Y cells, we found significant alteration in miRNA expression in Lamin A KO cells as compared to control cells (**Figure 3H**). Of note, the top upregulated miRNA species are members of the miRNA-17/92 cluster, one of the best-described oncogenic miRNA families [10,80]. In neuroblastoma, miRNA-17/92 expression is positively correlated with higher malignancy and poor overall outcome [33–34]. Moreover, high expression of the cluster is triggered by the primary neuroblastoma-associated oncogene, MYCN, and directly linked to enhanced cellular proliferation and impaired differentiation [33–34]. Hence, the miRNA deregulation found upon Lamin A KO in SHSY5Y contributes to the observed phenotypical changes, underlying the importance of precise regulation of RNAi activity in cancer progression.

Since RNAi is an essential cellular process, its activity is fine-tuned by various molecular mechanisms. RBPs are important factors that regulate the activity of RNAi predominately by acting as competitors, either against RISC components or competitors for target mRNA binding [32–34]. Most RBPs were shown to enhance RNAi activity by rearranging the secondary structure of target transcripts and thus improving miRNA binding accessibility [81]. Yet, a few RBPs were found to selectively inhibit miRNA-dependent destabilization of certain transcripts [32–34,82]. For instance, the ELAVL gene family encodes several RBPs responsible for stabilizing transcripts promoting cell proliferation by masking miRNA binding sites [32–33]. In our study, proteomics of AGO interactome from the nuclear fraction unveiled an interaction between AGO and FAM120A in both A375 and SHSY5Y cells (**Figure 5A,B**). This interaction was previously shown in stem cells and T47D breast cancer cells [65,67]. Of note, both studies used Flag-AGO2 overexpression systems [65,67], while we detected an interaction between endogenous AGO2 and FAM120A proteins of Lamin A KO cells (**Figure 5E,F**, **Figure 5K,L**). iCLIP of Flag-FAM120A and Flag-AGO2 in stem cells revealed a fraction of mRNAs that was bound by AGO2 and FAM120A simultaneously [65]. On those subsets of genes, AGO2 did not mediate target degradation [65]. Similarly, we found a profound overlap between transcripts bound by AGO proteins and FAM120A in the Lamin A KO cells (**Figure 6F,G, Supplemental Figure 6H,I**). Moreover, our data confirm that those targets common between AGO and FAM120A were stabilized (**Figure 6H,I, Supplemental Figure 6J,K**). The protective effect of FAM120A was found in both the cytoplasmic and nuclear fraction, suggesting that the direct interaction between AGO and FAM120A is dispensable for transcript binding and stabilization.

In this study, we revealed a new layer of regulation of RNAi. We uncovered that the downregulation of Lamin A in cancer cells, an alteration promoting cancer aggressiveness, is sufficient to trigger AGO2 nuclear influx accompanied by profound impairment of RNAi in both the cytoplasm and nucleus of Lamin A KO cells. In contrast, potent nuclear RNAi was observed in mESC, where nuclear AGO2 is expressed at high levels in steady-state conditions [18]. Lamin A loss triggered significant phenotypical alterations, including enhanced proliferation and dedifferentiation. Notably, we found that FAM120A protein co-binds AGO targets, rendering them stabilized, in Lamin A KO conditions. Corresponding observations were previously made in embryonic stem cells, albite to a lesser degree. Together, our data gives insights into the molecular mechanisms of fine-tuning RNAi activity in cancer cells.

## DATA AVAILABILITY

The mass spectrometry proteomics data have been deposited to the ProteomeXchange Consortium (http://proteomecentral.proteomexchange.org) via the PRIDE partner repository [83]. with the dataset identifier PXD042899. fPAR-CLIP sequencing data are available on the NCBI Short-Read Archive (SRA) under the accession number GSE261593. RNA sequencing data are available on the NCBI Short-Read Archive (SRA) under the accession number GSE235156. miRNA sequencing data are available on the NCBI Short-Read Archive (SRA) under the accession number GSE261422.

## SUPPLEMENTARY DATA

Supplementary Data are available at NAR online.

## AUTHOR CONTRIBUTIONS

Vivian Lobo: Formal analysis, Methodology, Writing—review & editing. Iwona Nowak: Formal analysis, Methodology, Writing—original draft. Carola Fernandez: Formal analysis, Methodology, Writing—review & editing. Ana Iris Correa Muler: Methodology, Writing—review & editing. Jakub O. Westholm: Formal analysis, Methodology, Writing—review & editing. Hsiang-Chi Huang: Methodology, Writing—review & editing. Ivo Fabrik: Formal analysis, Writing—review & editing. Hang T. Huynh: Methodology, Writing—review & editing. Evgeniia Shcherbinina: Methodology, Writing—review & editing. Melis Kanik: Methodology, Writing— review & editing. Anetta Härtlova: Formal analysis, Writing—review & editing. Daniel Benhalevy: Formal analysis, Writing—review & editing. Davide Angeletti: Formal analysis, Methodology, Writing—review & editing. Aishe A. Sarshad: Conceptualization, Funding acquisition, Supervision, Formal analysis, Methodology, Writing—original draft.

## ACKNOWLEDGEMENTS

The authors would like to acknowledge Dr. Ruth Palmer and Dr. Bengt Hallberg (University of Gothenburg) for the STE01 and CUT01 cancer cell lines, Maria Falkenberg (University of Gothenburg) for HAP1 cells. Volkan Sayin (University of Gothenburg) for providing us with Menadione. Dimitrios Anastasakis (NIIAMS, NIH) for technical support. Reyhaneh Rezaei for technical support. Sequencing was performed by the SNP&SEQ Technology Platform in Uppsala. The facility is part of the National Genomics Infrastructure (NGI) Sweden and Science for Life Laboratory. The SNP&SEQ Platform is also supported by the Swedish Research Council and the Knut and Alice Wallenberg Foundation. We acknowledge the Center for Cellular Imaging at the University of Gothenburg and the National Microscopy Infrastructure, NMI (VR-RFI 2019-00217) for providing assistance in microscopy. The proteomic analysis was performed at the Proteomics Core Facility at the University of Gothenburg, Sweden. The computations were enabled by resources in projects SNIC 2022/23-474 and SNIC 2022/22-996 provided by the National Academic Infrastructure for Supercomputing in Sweden (NAISS) at UPPMAX, funded by the Swedish Research Council through grant agreement no. 2022-06725.

