## Supplementary material for "Loss of Lamin A leads to the nuclear translocation of AGO2 and compromised RNA interference": Sup data

**Supplemental information**

**Supplemental Figure 1 related to Figure 1 (A)** Representative images of Lamin A/C and Lamin B1 immunoblots from A375 and SHSY5Y cells transfected with siRNAs against LMNA and LMNB1 respectively. GAPDH was used as the refence. Representative images of AGO2 immunoblots from cytoplasmic and nuclear fractions in cells transfected with the siRNAs for LMNA in **(B)** HEK293 and **(C)** U2OS cells. For biochemical fractionation experiments CALR served as cytoplasmic marker while LMNA and LMNB as nuclear markers. C, cytoplasmic fraction; N, nuclear fraction; CALR, Calreticulin; LMNA, Lamin A/C; Cont, cells treated with Lipofectamine RNAiMAX transfection agent; Scr, cells transfected with scrambled oligonucleotide; si*LMNA*, cells transfected with siRNAs against *LMNA*; si*LMNB1,* cells transfected with siRNAs against *LMNB1.*

**Supplemental Figure 2 related to Figure 2**. **(A)** Graphical representation of the CRISPR-Cas9 strategy used in the study. Numbers and arrows illustrate the location and direction of primers used in PCR experiments. PCR amplification of antibiotic resistance/EGFP cassette in (**B**) A375, (**C**) SHSY5Y and (**D**) HeLa Lamin KO cells. Primer numbers are indicated. (**E**) Bright field microscopy images of A375 (left panel) and SHSY5Y (right panel) Lamin A KO (upper panel) and WT (bottom panel) cells. Immunofluorescence staining of Lamin A, LMNB1, Phalloidin, Pol II, H3, GAPDH and DAPI in **(F)** A375 and **(G)** SHSY5Y WT and Lamin A KO cells. (**H**) Representative images of AGO2 immunoblots from cytoplasmic and nuclear fractions in HeLa WT and Lamin KO cells. Representative images of AGO1 and 3 immunoblots from cytoplasmic and nuclear fractions in **(I)** A375 and **(J)** SHSY5Y WT and Lamin KO cells. For biochemical fractionation experiments CALR served as cytoplasmic marker while LMNB1 served as nuclear markers. AGO2 expression levels in Lamin A KO vs WT **(K)** A375 and **(L)** SHSY5Y cells determined using RT-qPCR and 18S rRNA as reference gene, values for WT cells set as 1. Results were calculated from three independent experiments, additionally performed in three technical triplicates, and presented as the mean ± standard deviation. Representative images of AGO2 immunoblots from **(M)** A375 and **(N)** SHSY5Y WT and Lamin A KO cells. CALR was used as a reference. WT, wild-type cells; Lamin A KO, Lamin A KO cells; C, cytoplasmic fraction; N, nuclear fraction; CALR, calreticulin; LMNB1, Lamin B1. *P<0.05.

**Supplemental Figure 3 related to Figure 3.** Relative viability of Lamin A KO vs WT in **(A)** A375 and **(B)** SHSY5Y cells assessed by MTT assay. Graph present average from three independent experiments, performed in technical triplicates, ± standard deviation. Representative histograms of cell cycle analysis by flow cytometry in WT and Lamin A KO growing **(C)** A375 and **(D)** SHSY5Y cells. PCA of three independent RNAseq experiments in **(E)** A375 and **(F)** SHSY5Y. **(G)** Biological process annotations provided GO for transcripts that were significantly downregulated and upregulated upon Lamin A KO in A375 and SHSY5Y cells. PCA, principal component analysis; WT, wild-type cells; KO, Lamin A KO cells; GO, Gene Ontology; BP, biological processes. *P<0.05, **P<0.01.

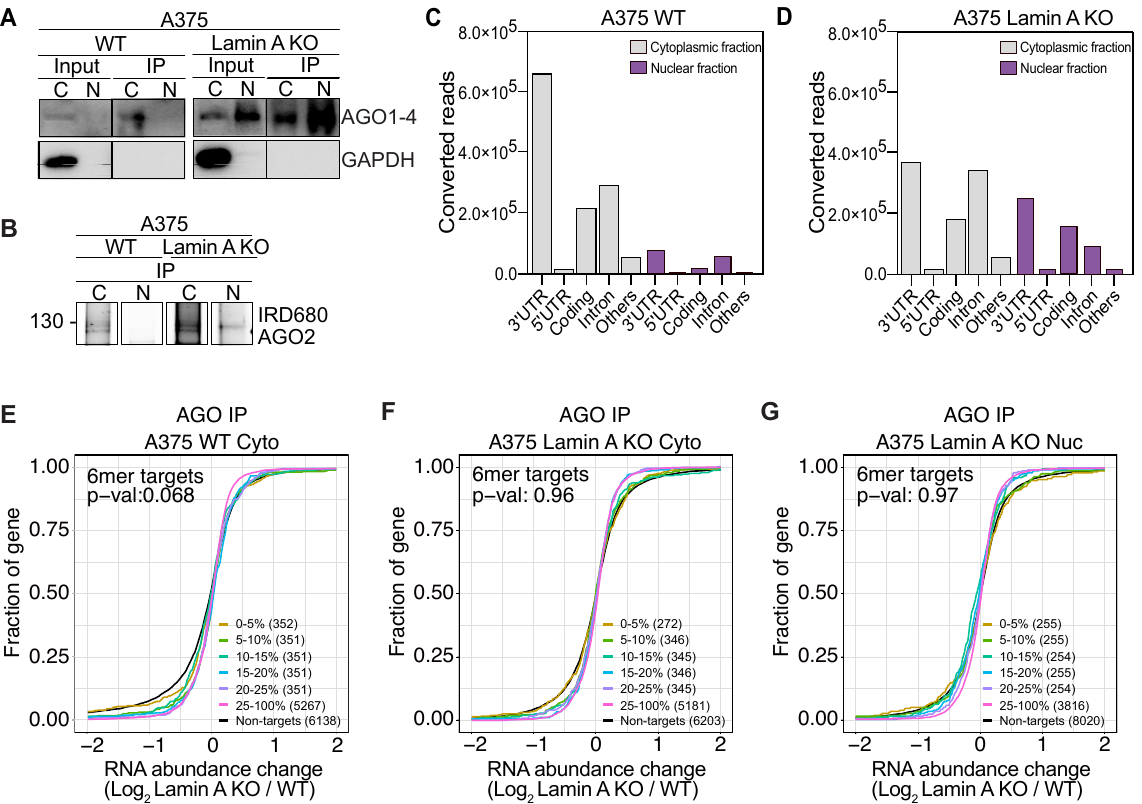

**Supplemental Figure 4 related to Figure 4. (A)** Representative images of AGO2 and GAPDH immunoblots from A375 WT and Lamin A KO cells after immunoprecipitation of AGO1-4 using the T6B peptide. (**B**) Fluorescence SDS-PAGE of 3’ labeled AGO1-4 RNP in WT and Lamin A KO A375 cells**.** Distribution of AGO fPAR-CLIP sequence reads across target RNAs in A375 **(C)** WT cytoplasmic and nuclear lysate and **(D)** Lamin A KO cytoplasmic and nuclear lysate. Cumulative distribution of abundance changes in RNA in A375 **(E)** WT cytoplasmic fraction **(F)** Lamin A KO cytoplasmic fraction and **(G)** Lamin A KO nuclear fraction. For cumulative distribution assays the targets were ranked by number of binding sites from top 5-20% and 25-100% and compared to non-targets. WT, wild-type cells, Lamin A KO, Lamin A knock out cells; C, cytoplasmic fraction; N, nuclear fraction; IP, immunoprecipitation; fPAR-CLIP, fluorescent photoactivatable ribonucleoside-enhanced crosslinking and immunoprecipitation; RNP, ribonucleoprotein complex.

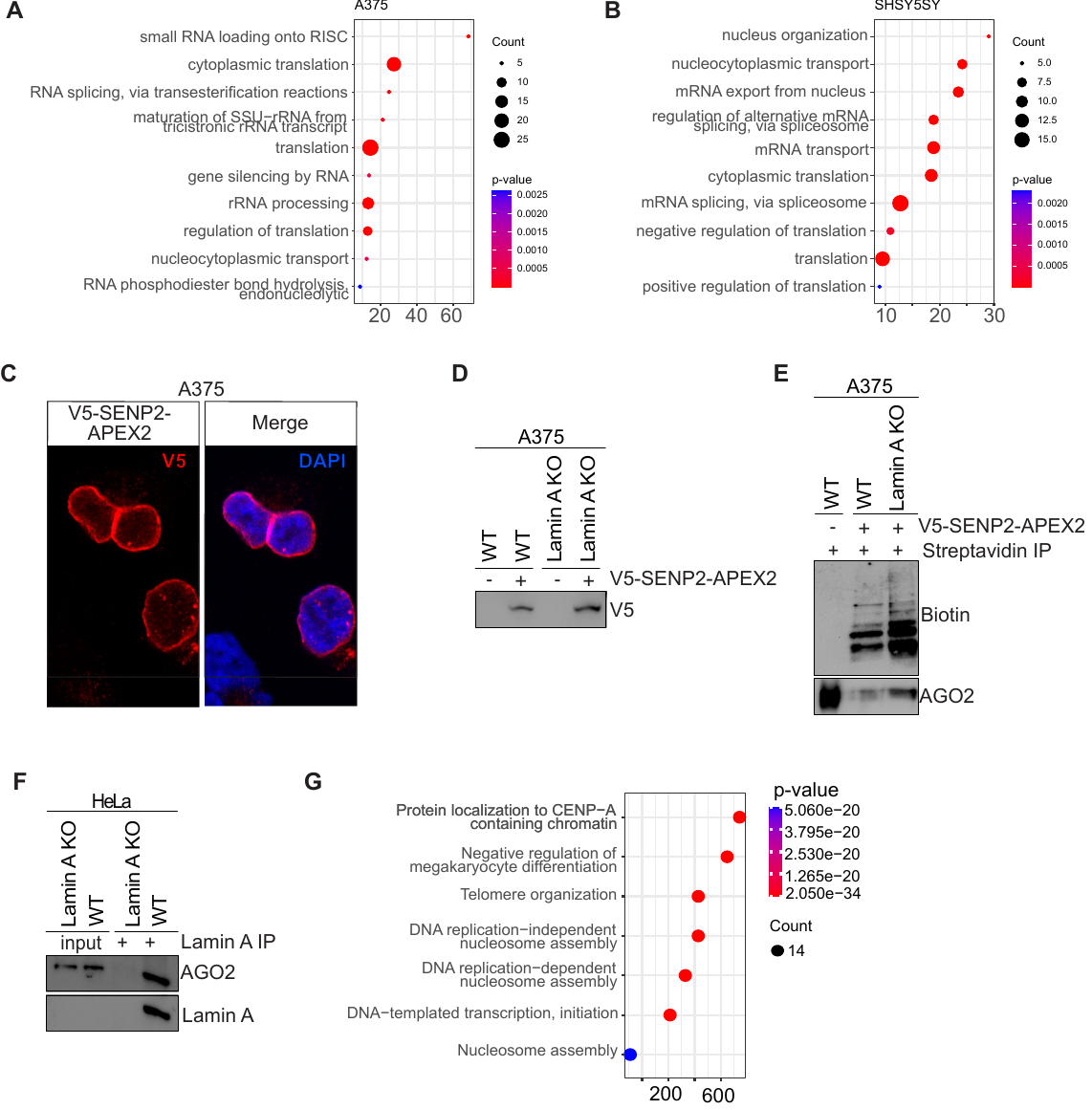

**Supplemental Figure 5 related to Figure 5.** Biological process annotations provided by GO for significantly upregulated processes the in Lamin A KO vs WT nuclear AGO IP mass spectrometry hits from **(A)** A375 and **(B)** SHSY5Y cells. **(C)** Immunofluorescence staining of V5-SENP2-APEX2 in A375 WT APEX-SENP2 and A375 Lamin A KO APEX2-SENP2 cells. **(D)** Representative image of V5 immunoblot of A375 WT APEX-SENP2 and A375 Lamin A KO APEX2-SENP2 cells as compared to parental cell lines. **(E)** Representative image of biotin and AGO2 immunoblot of streptavidin affinity purification from A375 WT APEX-SENP2 and A375 Lamin A KO APEX2-SENP2 cells as compared to parental cell line A375 WT. **(F)** Representative images of AGO2 and Lamin A immunoblots of co-immunoprecipitation assay in HeLa WT and Lamin A KO cells. **(G)** Biological process annotations provided by the Gene Ontology for significantly upregulated in Lamin A IP from WT vs Lamin A KO HeLa cells. WT, wild-type cells; Lamin A KO, Lamin A knock out cells; V5-SENP2-APEX2, cells expressing V5-SENP2-APEX2 protein; GO, Gene Ontology.

**Supplemental Figure 6 related to Figure 6.** Optimization of Menadione concentration in **(A)** A375 and **(B)** SHSY5Y WT cells. Assessment of H_2_O_2_ species in the media of Lamin A KO and WT **(C)** A375 and **(D)** SHSY5Y cells transfected with either GFP or LMNA expression plasmids after treatment with ROS-inducer (Menadione). Graph present average of three independent experiments, performed in technical triplicates, mean ± standard deviation. **(E)** Representative images of FAM102A and PCNA immunoblots from A375 WT and Lamin A KO cells before and after immunoprecipitation of FAM120A. **(F)** Fluorescence SDS-PAGE of 3’ labeled FAM120A RNP in WT and Lamin A KO A375 cells. **(G)** Distribution of FAM120A fPAR-CLIP sequence reads across target RNAs in A375 Lamin A KO cytoplasmic and nuclear lysate. Overlap between AGO and FAM120A fPAR-CLIP targets in A375 cells from **(H)** cytoplasmic fraction and **(I)** nuclear fraction. Cumulative distribution of abundance changes in RNA of AGO:FAM120A overlapping targets in A375 Lamin A KO **(J)** cytoplasmic fraction and **(K)** nuclear fraction. For cumulative distribution assays the targets were ranked by 25 percentile and compared to non-targets. WT, wild-type cells, Lamin A KO, Lamin A knock out cells; C, cytoplasmic fraction; N, nuclear fraction; IP, immunoprecipitation; fPAR-CLIP, fluorescent photoactivatable ribonucleoside-enhanced crosslinking and immunoprecipitation; RNP, ribonucleoprotein complex. **P<0.01,***P<0.001,****p<0.0001.
